# Ly6C defines a subset of memory-like CD27^+^ γδ T cells with inducible cancer-killing function

**DOI:** 10.1101/2020.09.08.287854

**Authors:** Robert Wiesheu, Sarah C. Edwards, Ann Hedley, Kristina Kirschner, Marie Tosolini, Jean-Jacques Fournie, Anna Kilbey, Sarah-Jane Remak, Crispin Miller, Karen Blyth, Seth B. Coffelt

## Abstract

In mice, IFNγ-producing γδ T cells that express the co-stimulatory molecule, CD27, play a critical role in host defence and anti-tumour immunity. However, their phenotypic diversity, composition in peripheral and secondary lymphoid organs, similarity to αβ T cells as well as homology with human γδ T cells is poorly understood. Here, using single cell RNA sequencing, we show that CD27^+^ γδ T cells consist of two major clusters, which are distinguished by expression of Ly6C. We demonstrate that CD27^+^Ly6C^—^ γδ T cells exhibit a naïve T cell-like phenotype, whereas CD27^+^Ly6C^+^ γδ T cells display a memory-like phenotype, produce several NK cell-related and cytotoxic molecules and are highly similar to both mouse CD8^+^ T cells and mature human γδ T cells. In a breast cancer mouse model, depletion of CD27^+^ γδ T cells failed to affect tumour growth, but these cells could be coerced into killing cancer cells after expansion *ex vivo*. These results identify novel subsets of γδ T cells in mice that are comparable to human γδ T cells, opening new opportunities for γδ T cell-based cancer immunotherapy research.

## INTRODUCTION

γδ T cells are a rare population of T cell receptor (TCR)-expressing lymphocytes that possess features of both innate and adaptive immune cells (Hayday, 2019). One of the first publications describing γδ T cells reported on their cancer-killing ability (Bank *et al*, 1986). Their importance in anti-tumour immunity was further corroborated in 2001 using a squamous cell carcinoma mouse model of skin cancer crossed with γδ T cell-deficient mice (Girardi *et al*, 2001). Since that time, several studies have identified γδ T cells as key players in cancer progression (Silva-Santos *et al*, 2019), and their abundance in human tumours can be used as a predictor of patient outcome (Gentles *et al*, 2015; Ma *et al*, 2012; Meraviglia *et al*, 2017; Wu *et al*, 2014). Consequently, efforts to exploit γδ T cells for cancer immunotherapy have garnered a great deal of attention (Sebestyen *et al*, 2020; Silva-Santos *et al.*, 2019). The anti-tumour properties of γδ T cells are distinctive from αβ T cells. In particular, their independence from MHC molecules makes them easily transferrable between cancer patients and ideal for low antigen-expressing or MHC-deficient tumours. For human clinical studies, the Vγ9Vδ2 and Vδ1 cell subsets have been adoptively transferred into cancer patients with some successes and some failures (Sebestyen *et al.*, 2020; Silva-Santos *et al.*, 2019). Researchers are currently testing new strategies to enhance cytotoxic function of the Vγ9Vδ2 and Vδ1 cell subsets for adoptive cell transfer with focus on *ex vivo* expansion protocols that maximize their killing capacity (Almeida *et al*, 2016; Di Lorenzo *et al*, 2019). Other approaches to specifically modify γδ T cells for cancer immunotherapy include the generation of γδ chimeric antigen receptor (CAR)-T cells (Mirzaei *et al*, 2016; Rischer *et al*, 2004) and T cells engineered with defined γδTCRs (TEGs) (Gründer *et al*, 2012; Marcu-Malina *et al*, 2011). However, mechanistic studies on the anti-tumour properties of these γδ T cell products and their influence on other immune cells in the tumour microenvironment (TME) are limited, because the appropriate immunocompetent mouse models are lacking.

In mice, γδ T cells consist of several subsets that include tissue resident – such as Vγ5 cells of the skin and Vγ7 cells of the intestine – and circulating cells. The circulating cells expressing Vγ1 or Vγ4 T cell receptor chains are divided into functionally distinct subgroups, largely based on their cytokine production and expression of the co-stimulatory molecule, CD27 (Ribot *et al*, 2009). The two major subgroups are CD27^+^ IFNγ-producing γδ T cells and CD27^—^ IL-17-producing γδ T cells. Most reports to date show opposing roles for these subgroups in the context of cancer. CD27^—^ IL-17-producing γδ T cells promote cancer progression and metastasis (Coffelt *et al*, 2015; Housseau *et al*, 2016; Ma *et al*, 2014; Patin *et al*, 2018; Rei *et al*, 2014; Van hede *et al*, 2017; Wakita *et al*, 2010), whereas CD27^+^ IFNγ-producing γδ T cells counteract cancer progression (Dadi *et al*, 2016; Gao *et al*, 2003; He *et al*, 2010; Lanca *et al*, 2013; Liu *et al*, 2008; Street *et al*, 2004). Mechanisms by which IFNγ-producing γδ T cells oppose tumour growth are poorly defined. For example, it is unclear which molecules are recognized by their TCRs or whether other receptors, such as NKG2D, are required for their activation. However, these cells can provide an early source of IFNγ that induces upregulation of MHC-class I expression and leads to the recruitment of cytotoxic CD8^+^ T cells into the tumour microenvironment (Gao *et al.*, 2003; Riond *et al*, 2009). CD27^+^ γδ T cells also use other cytotoxic molecules to kill cancer cells, such as granzyme B (GzmB) and perforin (Dadi *et al.*, 2016). In addition to the studies focused on the endogenous anti-tumour role of CD27^+^ IFNγ-producing γδ T cells, a few studies have explored their ability to control tumour growth after *ex vivo* expansion and adoptive cell transfer into tumour-bearing mice (Beck *et al*, 2010; Cao *et al*, 2016b; He *et al.*, 2010; Liu *et al.*, 2008; Street *et al.*, 2004). Mirroring the outcomes of experiments using human Vγ9Vδ2 or Vδ1 cells, these reports indicate that expanded γδ T cells from mice are capable of thwarting tumour growth. Despite these efforts, however, the degree of homology between human and mouse γδ T cell subsets is largely unknown, dampening the enthusiasm for the development of syngeneic tumour models that utilise mouse γδ T cell products. Therefore, we set out to characterize the phenotype and function of anti-tumour γδ T cells in mice and their relationship to human γδ T cells.

Here, we provide insight into the nature of mouse CD27^+^ γδ T cells using single cell RNA sequencing. We identify Ly6C as a marker delineating two populations: CD27^+^Ly6C^+^ γδ T cells and CD27^+^Ly6C^—^ γδ T cells. CD27^+^Ly6C^—^ γδ T cells express lower levels of cytotoxic molecules, NK cell-related molecules and more naïve T cell-like markers when compared to CD27^+^Ly6C^+^ γδ T cells, which exhibit an effector/memory-like phenotype. In relation to human γδ T cells, the gene expression signature of mouse CD27^+^Ly6C^—^ γδ T cells aligns with the transcriptome of human naïve T cells, while the gene expression signature of mouse CD27^+^Ly6C^+^ γδ T cells aligns with the transcriptome of mature human γδ T cells, CD8^+^ T cells and NK cells. Because CD27^+^ γδ T cells consisted mainly of Vγ1 cells, we depleted Vγ1 cells in a syngeneic breast cancer mouse model to assess their effects on tumour growth. Loss of endogenous Vγ1 cells failed to influence cancer progression, but their cancer killing ability could be induced by *ex vivo* expansion with IL-15, T cell receptor stimulation through CD3 and co-stimulation through CD28. These data offer insight into the phenotype and activation status of anti-tumour γδ T cell subsets in mice, whose utility may be exploited to better understand human γδ T cell-based cancer immunotherapies.

## RESULTS

### Lung CD27^+^ γδ T cells cluster into three major groups

We previously defined the pro-metastatic role of IL-17-producing (CD27^—^) γδ T cells in the lungs of mammary tumour-bearing mice (Coffelt *et al.*, 2015; Wellenstein *et al*, 2019). To better understand the distinctive phenotype and heterogeneity of CD27^+^ γδ T cells in this organ, we sorted total γδ T cells from the lungs of wild-type (WT) mice and analysed these cells by single cell RNA sequencing (scRNAseq) using the Chromium 10X platform. Cells with *Cd27* expression were computationally separated for further downstream analysis, resulting in 458 CD27^+^ γδ T cells. t-Distributed Stochastic Neighbour Embedding (t-SNE) was utilised for visualization of the data, which identified three distinct clusters of CD27^+^ γδ T cells, with Cluster 2 being the most transcriptionally different from Clusters 0 and 1 (**Fig 1A**). We then interrogated the gene expression differences between the three clusters and it became evident why Cluster 2 separated from Clusters 0 and 1 (**Fig 1B**). The top differentially expressed genes of Cluster 2 shared a highly similar gene signature (*Cd163l1, Cxcr6, Bcl2a1b, Lgals3, Tmem176a/b, S100a4*) with recently reported IL-17-producing γδ T cell populations in the skin, lymph nodes (LNs) and adipose tissue (Chen *et al*, 2019; Kohlgruber *et al*, 2018; Tan *et al*, 2019). The *Cd163l1* gene encodes the SCART1 protein that is used as a surrogate marker for Vγ6 cells with high capacity to produce IL-17A (Kisielow *et al*, 2008; Tan *et al.*, 2019), indicating that Cluster 2 is most likely a lung-resident γδ T cell subset that expresses high transcript levels of *Cd27* mRNA but lacks CD27 protein expression on the cell surface. Since our research question pertained to defining novel subsets of *bona fide* CD27^+^ γδ T cells our subsequent efforts for the remainder of the study were solely focused on Clusters 0 and 1.

**Figure 1.**
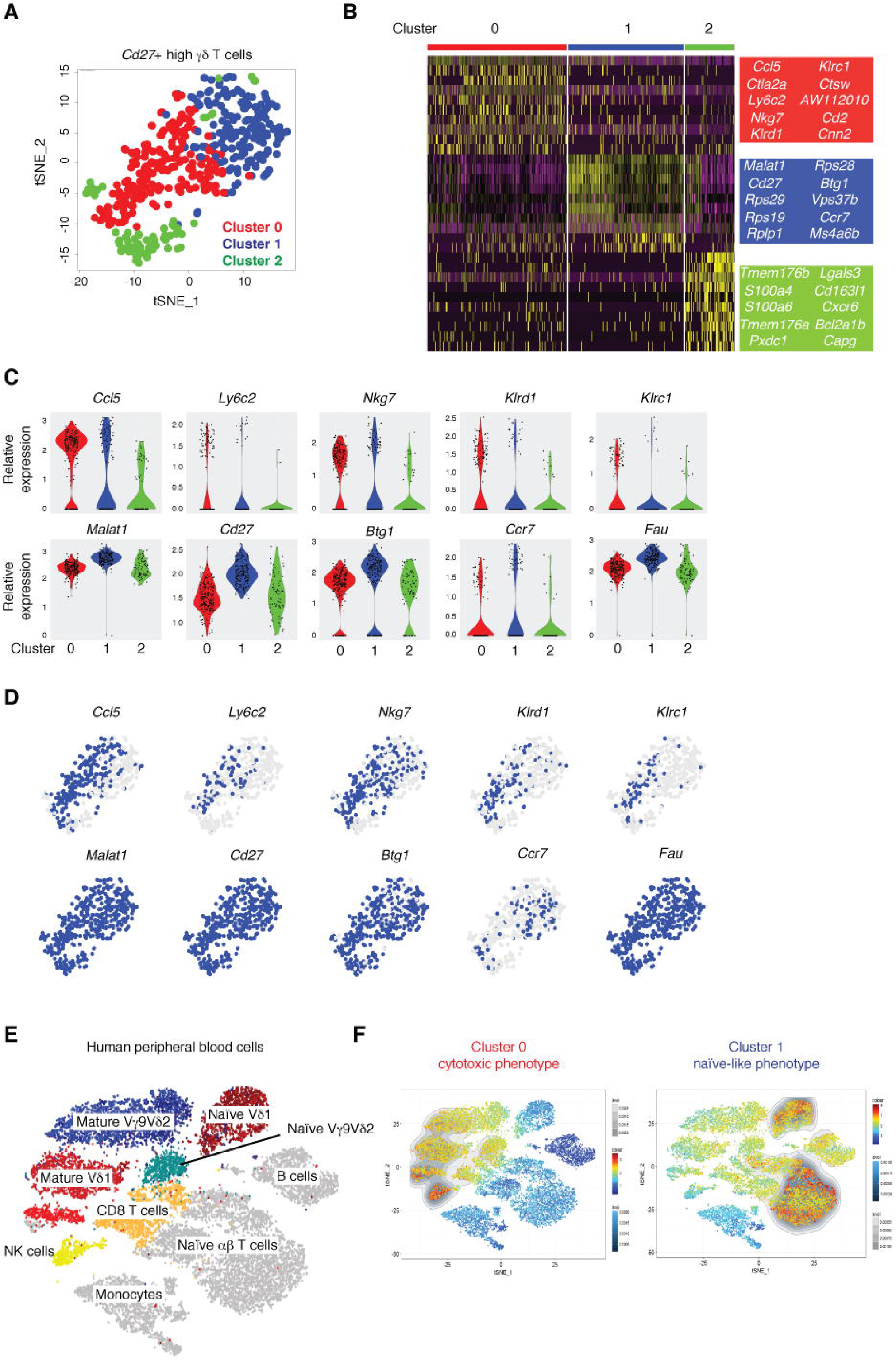
Single-cell RNA sequencing (scRNAseq) reveals distinct populations of mouse CD27^+^ γδ T cells with similarities to human lymphocytes. γδ T cells from lungs of two wild-type (WT) FVB/n mice were sorted and analyzed by scRNAseq on the Chromium 10X platform in two separate runs. **(A)** tSNE visualization of 458 individual lung CD27^+^ γδ T cells (n = 4 mice), colour-coded by cluster. **(B)** Heatmap of the top 10 genes from each of the 3 clusters identified in A, where each column represents the gene expression profile of a single cell. Gene expression is colour-coded with a scale based on z-score distribution, from low (purple) to high (yellow). **(C)** Violin plots showing expression levels of selected genes from the clusters identified in A. **(D)** Feature plots of the same genes shown in C, depicting expression levels by cell. Blue indicates high expression and grey indicates no expression. **(E)** tSNE visualization of a human scRNAseq dataset containing ~10^4^ PBMCs and γδ T cells from 3 individual healthy donors. Clusters are coded by different colours and labelled by cell type. **(F)** Gene signatures from Cluster 0 (left panel) and Cluster 1 (right panel) displayed on the tSNE map from E by Single-Cell Signature Viewer. The coloured scales represent the degree of transcriptional similarity where red indicates high similarity and dark blue indicates low similarity. The grey scale represents density distribution of similarity scores.

Cells from Cluster 0 were enriched in NK cell-associated genes (*Cd160*, *Ncr1*, *Nkg7, Klrd1, Klrc1, Gzma*) as well as the cytotoxic markers Cathepsin W and cytotoxic T lymphocyte-associated protein 2 complex (*Ctsw* and *Ctla2a*). Additionally, cells from Cluster 0 were enriched for the chemoattractant, *Ccl5, Ifng* and *Ly6c2* (**Fig 1B-D, Table 1**). The *Ly6c2* gene encodes the Ly6C protein, which is a cell differentiation antigen commonly used to identify cells of the myeloid compartment in mice, particularly monocytes and neutrophils. In Cluster 1, we found that *Malat1, Btg1, Ccr7, S1pr1* and *Fau* as well as many ribosomal proteins (*Rps1*, *Rps19*, *Rps28* and *Rps29*) are expressed to higher degree than in the cells from Cluster 0 (**Fig 1B-D, Table 2**). *Ccr7* expression was unique among these genes, since its expression was largely restricted to Cluster 1, whereas only subtle differences in expression of *Malat1*, *Btg1* and *Fau* were observed between Clusters 0 and 1 (**Fig 1B-D**). Naïve CD4^+^ T cells produce high levels of metastasis-associated lung adenocarcinoma transcript 1 (MALAT1), and MALAT1 is immediately downregulated upon activation of these cells (Hewitson *et al*, 2020). Similarly, the transcription factor BTG1 plays a crucial role in the maintenance of a T cell resting state, better known as T cell quiescence, prior to T cell activation (Hwang *et al*, 2020). In humans, the tumour-suppressor FAU interacts with the protein BCL-G to regulate apoptosis (Pickard *et al*, 2011). CCR7 is a vital lymph node homing receptor expressed by naïve T cells (Girard *et al*, 2012; Stein *et al*, 2000). S1PR1 is critical for exit of naïve T cells from lymph nodes (Baeyens *et al*, 2015). Taken together, the transcriptomic profile of CD27^+^ γδ T cells in Cluster 1 is akin to naïve T cells, whereas cells in Cluster 0 are enriched in cytotoxicity-associated gene signatures.

**Table 1.**
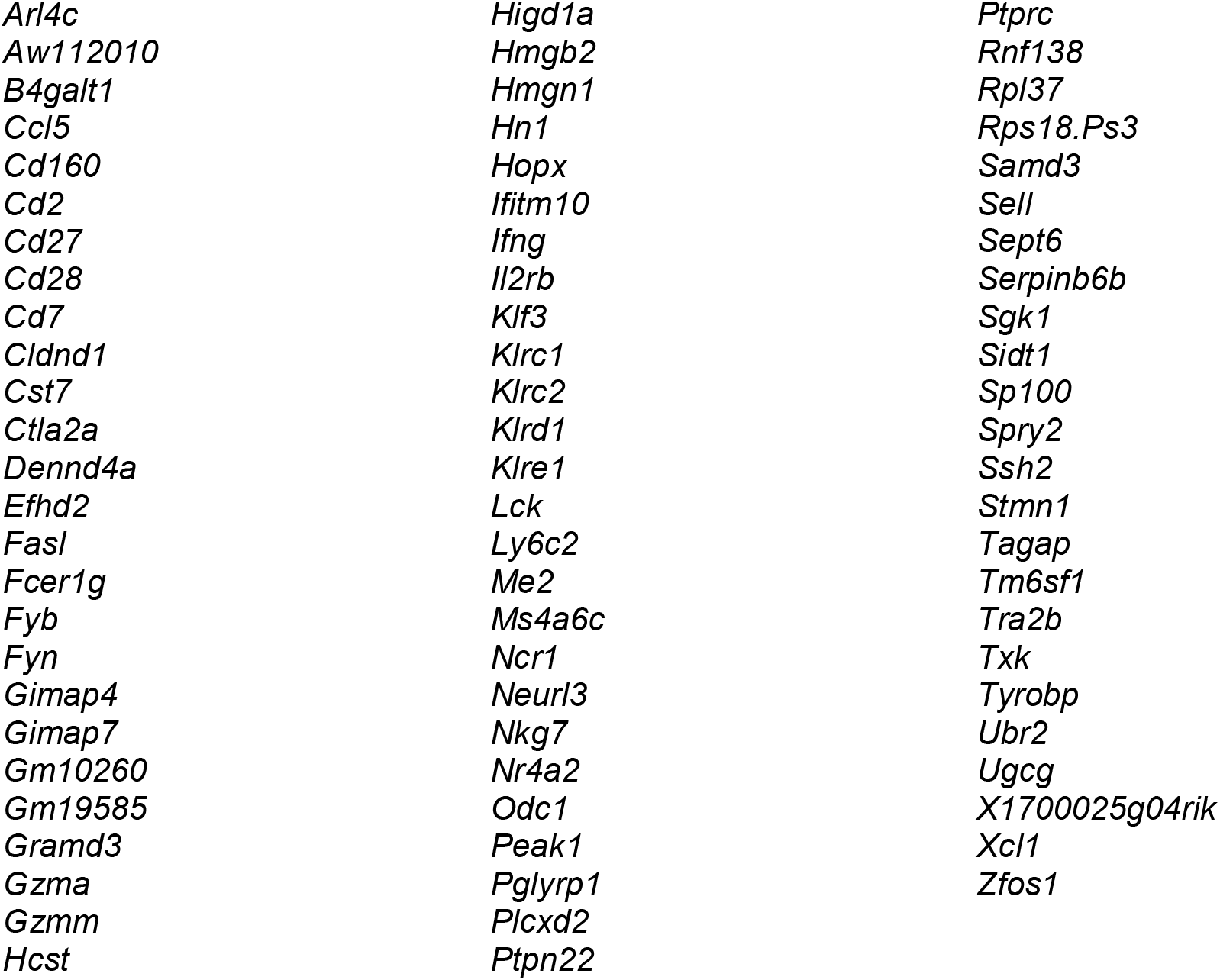
Genes enriched in cells from Cluster 0.

**Table 2.**
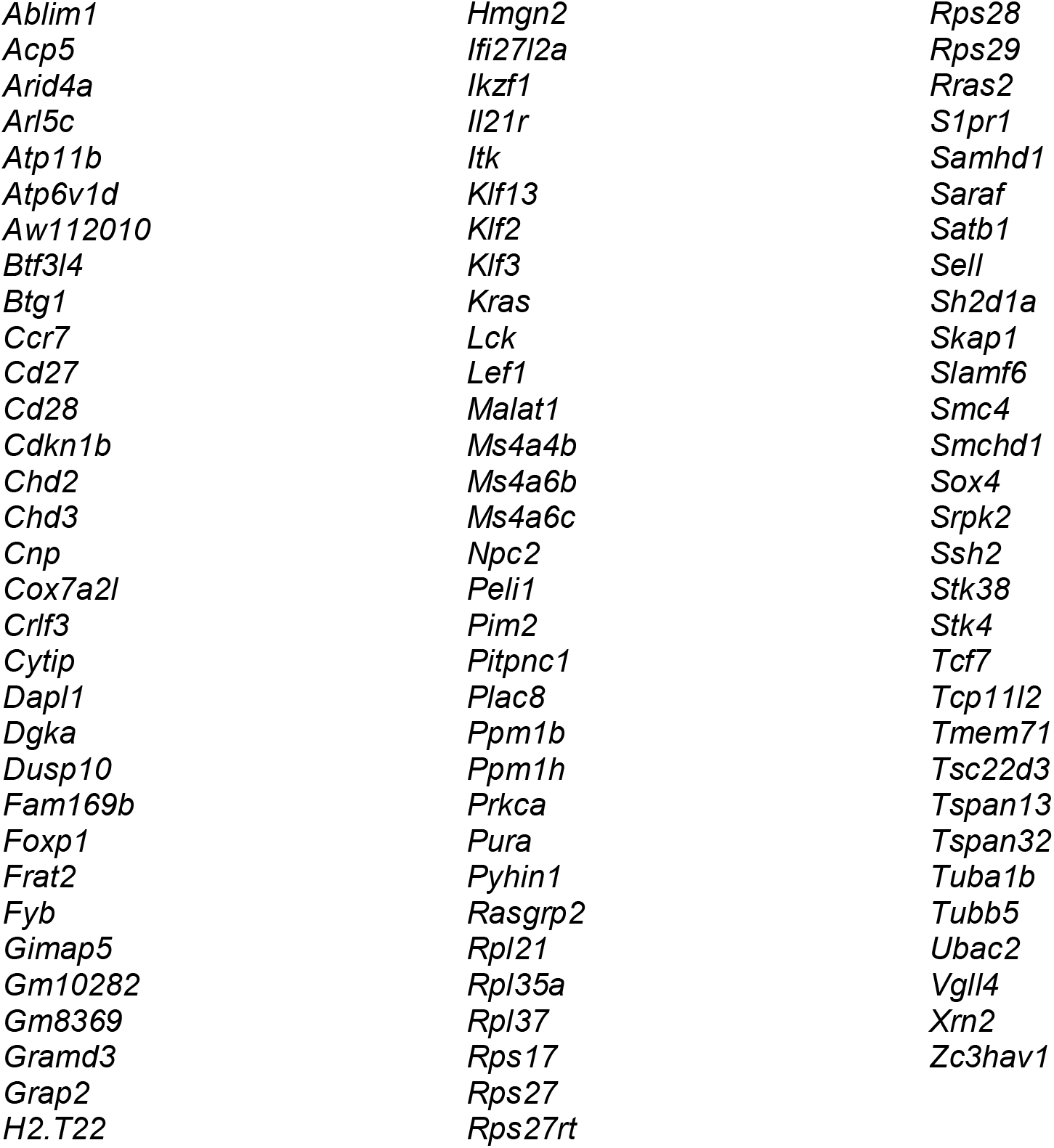
Genes enriched in cells from Cluster 1.

### Mouse γδ T cell transcriptional signatures align with human γδ T cells

The degree of homology between mouse and human γδ T cells is controversial and poorly understood. Therefore, we investigated the transcriptomic similarity between CD27^+^ γδ T cells from mice and human γδ T cells. We generated gene signatures from the scRNAseq data shown in Fig 1A-D for the cytotoxic group of cells in Cluster 0 and the naïve-like cells in Cluster 1, consisting of 76 and 94 genes, respectively (**Tables 1, 2**). These gene signatures were compared to a publicly available human scRNAseq data from approximately 10,000 cells comprising Vγ9Vδ2 cells, Vδ1 cells, CD8^+^ T cells, CD4^+^ T cells, NK cells, B cells and monocytes purified from three healthy donors (Pizzolato *et al*, 2019) (**Fig 1E**). Single-Cell_Signature_Explorer methodology was used to compare mouse and human data (Pont *et al*, 2019). When projected across the human dataset, the gene signatures of the two murine CD27^+^ γδ T cell clusters corresponded to well-defined human cell clusters. The gene signature from Cluster 0, which was enriched in cytotoxic molecules and *Ly6c2* (**Fig 1B-D**), mapped onto mature, terminally differentiated Vγ9Vδ2 cells, Vδ1 cells, NK cells and CD8^+^ T cells (**Fig 1F**). By contrast, the gene signature from Cluster 1, which contained more T cell naïve-like genes, such as *Ccr7* and *S1pr1* (**Fig 1B-D**), mapped onto naïve Vγ9Vδ2 cells, Vδ1 cells and αβ T cells (**Fig 1F**). These data not only underscore the high degree of similarity of γδ T cells between species, but they also provide a strong rationale to investigate the function of anti-tumour γδ T cells from mice to inform human γδ T cell biology.

### Ly6C defines a subset of CD27^+^ γδ T cells with a cytotoxic phenotype

Having identified two transcriptionally distinct subsets of CD27^+^ γδ T cells in mouse lung tissue by scRNAseq (**Fig 1A-D**), we next determined whether these two subsets could be distinguished by protein-based assays. We chose Ly6C and CCR7 as representative markers for Cluster 0 and Cluster 1, respectively, because of the robust and opposing expression levels observed by scRNAseq, as well as their location on the cell surface. Ly6C is a distinguishing marker of mouse monocytes and neutrophils. Ly6C is also expressed on memory CD8^+^ T cells, which exhibit increased IFNγ and granzyme B production (Allam *et al*, 2009; Marshall *et al*, 2011). In addition, Ly6C together with the activation marker CD44 defines γδ T cell subsets in the lymph nodes (Lombes *et al*, 2015); although, it is unknown whether γδ T cells maintain expression of Ly6C outside secondary lymphoid organs. CCR7 is a well-studied chemokine receptor critical for trafficking both T cells and dendritic cells (Girard *et al.*, 2012; Randolph, 2016; Roberts *et al*, 2016; Stein *et al.*, 2000). We used multi-parameter flow cytometry to analyse the expression of Ly6C and CCR7 on CD27^+^ γδ T cells from WT mouse lung, and extended this analysis to lymph nodes (LN) and spleen of naïve mice. This analysis revealed that approximately 30% of CD27^+^ γδ T cells express Ly6C across all tissues examined (**Fig 2A-B**). This uniformity was not seen with CCR7 expression. Expression of CCR7 was highest on lung CD27^+^ γδ T cells and lower on CD27^+^ γδ T cells from spleen and LN (**Fig 2A-B**), suggesting that this chemokine receptor is down-regulated in secondary lymphoid organs once CD27^+^ γδ T cells home to these sites. Because CCR7 expression patterns differed between tissues, we focused on Ly6C as a marker that may globally segregate the phenotypically distinct CD27^+^ γδ T cell clusters observed in scRNAseq analysis.

**Figure 2.**
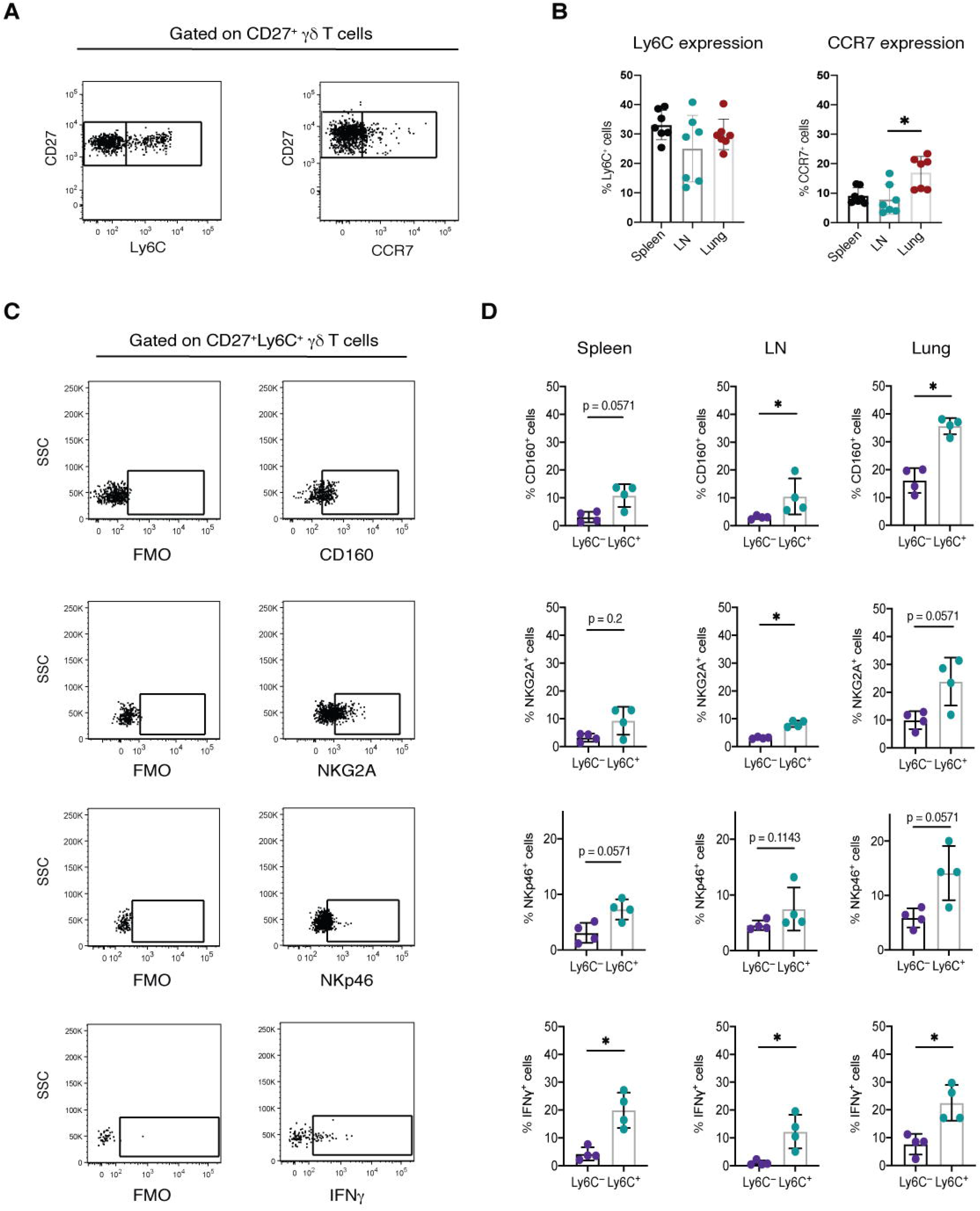
Ly6C defines a subset of cytotoxic CD27^+^ γδ T cells. Single-cell suspensions of spleen, lymph node (LN) and lung from WT mice were analyzed by flow cytometry. **(A)** Representative dot plots of Ly6C and CCR7 staining. Viable single cells were gated on CD3^+^ and TCRδ^+^ cells, followed by CD27^+^ cells. **(B)** Bar graphs showing the proportion of Ly6C^+^ cells among CD27^+^ γδ T cells in spleen, LN and lung (n = 7/group). Each dot represents one mouse. Data are represented as mean ± SD. \**p* < 0.05 as determined by one-way ANOVA followed by Dunn’s posthoc test. **(C)** Representative dot plots of CD160, NKG2A, NKp46 and IFNγ staining, after gating on CD3^+^TCRδ^+^ cells followed CD27^+^Ly6C^+^ cells. Fluorescence minus one (FMO) controls were used to draw gates. SSC = side scatter. **(D)** Bar graphs showing the proportion of CD160, NKG2A, NKp46 or IFNγ positive cells for CD27^+^Ly6C^—^ and CD27^+^Ly6C^+^ γδ T cell populations (n = 4/group) in spleen, LN and lung. Each dot represents one mouse. Data are represented as mean ± SD. \**p* < 0.05 as determined by Mann-Whitney U test.

The scRNAseq data indicated that CD27^+^ γδ T cells expressing Ly6C are more cytotoxic in nature than cells lacking Ly6C expression (**Fig 1A-D**). To validate this observation at the protein level, we chose four molecules from the gene signature list of Cluster 0 (**Table 1**) for which antibodies were available: CD160, NKG2A (the product of the *Klrc1* gene), NKp46 (the product of the *Ncr1* gene) and IFNγ. CD160 functions as a stimulatory molecule on NK cells and it induces IFNγ (Tu *et al*, 2015). NKG2A is an inhibitory molecule that regulates NK cell activation (Andre *et al*, 2018). NKp46 is a common marker for NK cells, whose activation stimulates NK cell cytolytic activity (Sivori *et al*, 1997). We analysed the expression of these molecules in CD27^+^Ly6C^—^ and CD27^+^Ly6C^+^ γδ T cells from spleen, LN and lung. We observed that CD27^+^Ly6C^+^ γδ T cells displayed higher expression levels of CD160, NKG2A and IFNγ regardless of tissue type, when compared with CD27^+^Ly6C^—^ γδ T cells (**Fig 2C-D**). NKp46 expression was higher on CD27^+^Ly6C^+^ γδ T cells than CD27^+^Ly6C^—^ γδ T cells in spleen and lung tissue. Overall, this phenotypic validation of scRNAseq data indicates that CD27^+^Ly6C^+^ γδ T cells (Cluster 0) represent a subset cells with increased cytotoxic ability, whereas CD27^+^Ly6C^—^ γδ T cells (Cluster 1) reside in a less activated or naïve state.

### Ly6C expression is enriched on memory/effector-like γδ T cells

Although γδ T cells are considered innate-like cells, the phenotype and function of CD27^+^ γδ T cells share certain aspects of adaptive immunity, including immunological memory that persists long-term (Lalor & McLoughlin, 2016). To this end, we measured naïve (CD62L^+^CD44^—^ cells), memory (CD62L^+^CD44^+^ cells) and effector memory (CD62L^—^CD44^+^ cells) status in total CD27^+^ γδ T cells from spleen, LN and lung of WT mice (**Fig 3A**). Naïve CD27^+^ γδ T cells were most abundant in LN, while spleen contained the highest proportion of memory CD27^+^ γδ T cells and lung exhibited the highest proportion of effector memory CD27^+^ γδ T cells (**Fig 3B**). We compared this distribution pattern with the proportions of naïve/memory/effector memory CD4^+^ and CD8^+^ T cells across the same organs. This comparison yielded a similar pattern of distribution, where LN contained the highest proportions of naïve CD4^+^ and CD8^+^ T cells and lowest proportions of effector memory CD4^+^ and CD8^+^ T cells (**EV1A**). The differences in proportions of naïve, memory and effector memory cell between secondary lymphoid organs and lung is consistent with T cell biology: lower numbers of antigen-inexperienced T cells in secondary organs and higher numbers of antigen-experienced T cells in peripheral organs.

**Figure 3.**
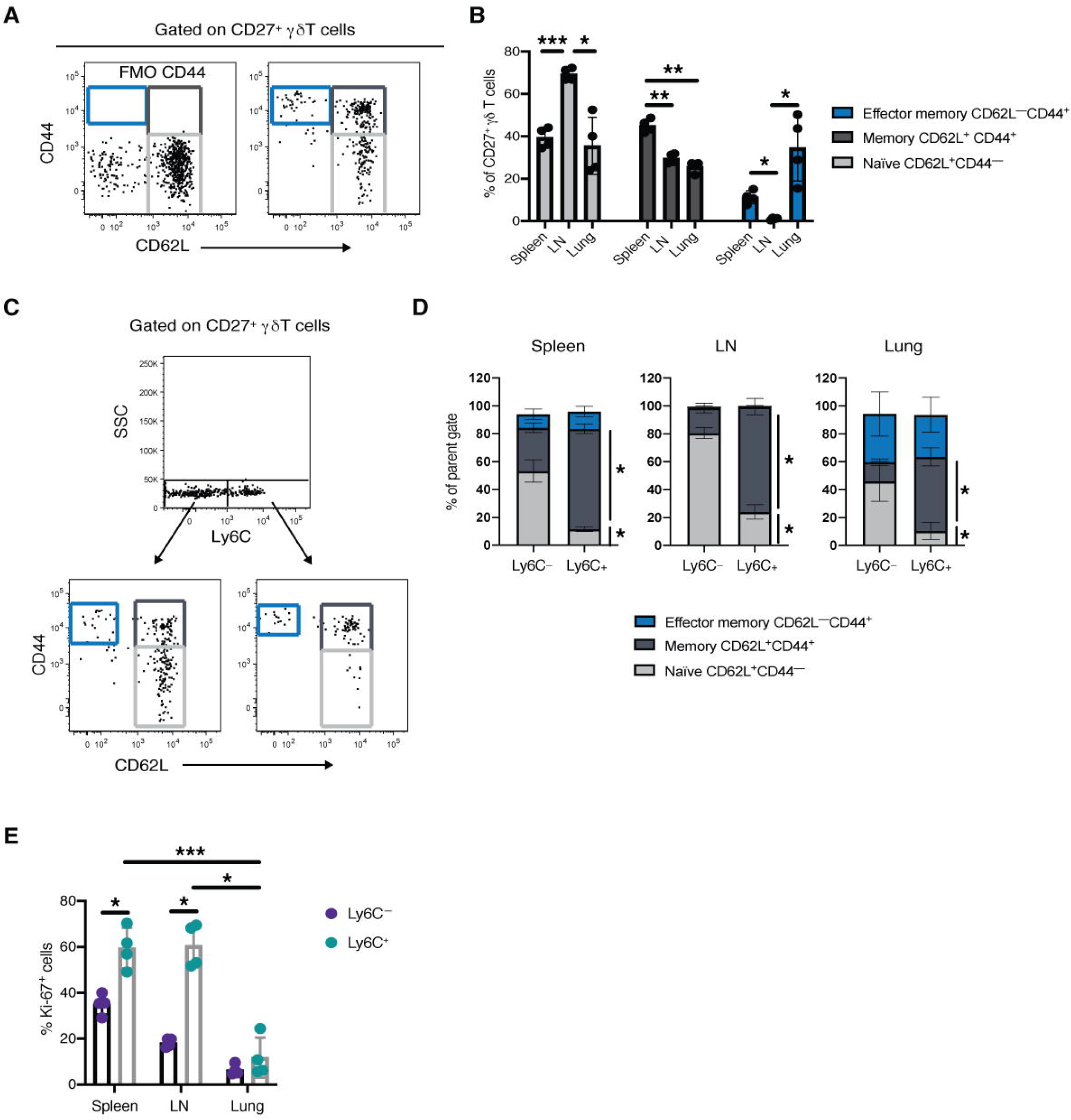
CD27^+^Ly6C^+^ γδ T cells display a memory/effector-like phenotype. Single-cell suspensions of spleen, lymph node (LN) and lung from WT mice were analyzed by flow cytometry. **(A)** Representative dot plots of CD44 and CD62L staining, after gating on CD3^+^TCRδ^+^ cells followed CD27^+^ cells. Fluorescence minus one (FMO) controls were used to draw gates. Blue box indicates effector memory cells; dark grey indicates memory cells; and light grey indicates naïve T cells. **(B)** Bar graphs showing the proportion of naïve T cells (CD62L^+^CD44^—^), memory (CD62L^+^CD44^+^) and effector memory (CD62L^—^CD44^+^) among CD27^+^ γδ T cells in spleen, LN and lung (n = 4/group). Each dot represents one mouse. Data are represented as mean ± SD. \**p* < 0.05, \*\**p* < 0.01, and \*\*\**p* < 0.001 as determined by repeated measures ANOVA followed by Tukey’s posthoc test. **(C)** Representative dot plots of CD3^+^TCRδ^+^CD27^+^ cells segregated into Ly6C^—^ and Ly6C^+^ subsets, followed by CD44 and CD62L staining. Blue boxes indicate effector memory cells; dark grey indicates memory cells; and light grey indicates naïve T cells. **(D)** Bar graphs showing the proportion of effector memory, memory and naïve T cells in CD27^+^Ly6C^—^ and CD27^+^Ly6C^+^ γδ T cells from spleen, LN and lung (n = 4/group). Data are represented as mean ± SD. \**p* < 0.05 as determined by Mann-Whitney U test. **(E)** Bar graphs showing the proportion of Ki67^+^ cells among CD27^+^Ly6C^—^ and CD27^+^Ly6C^+^ γδ T cells from spleen, LN and lung (n = 4/group). Each dot represents one mouse. Data are represented as mean ± SD. \**p* < 0.05 and \*\*\**p* < 0.001 as determined by Mann-Whitney U test or repeated measures ANOVA followed by Tukey’s posthoc test.

Based on the scRNAseq analysis and phenotypic data (**Fig 1–2**), which revealed an association between CD27^+^Ly6C^+^ γδ T cells and a cytotoxic, effector phenotype, we hypothesized that Ly6C is a surrogate marker for effector memory CD27^+^ γδ T cells. To address this hypothesis, we analysed CD27^+^Ly6C^—^ γδ T cells versus CD27^+^Ly6C^+^ γδ T cells from spleen, LN and lung of WT mice using the gating strategy shown in **Fig 3C**. This analysis showed that approximately 50% or more of CD27^+^Ly6C^—^ γδ T cells consist of CD62L^+^CD44^—^ naïve-like cells in the three tissues examined. The remainder of the CD27^+^Ly6C^—^ γδ T cells expressed CD44, associating them with memory-like cells (**Fig 3D**). By contrast, nearly all CD27^+^Ly6C^+^ γδ T cells (~80% or more) expressed CD44 in every tissue type, associating them with memory- and effector memory-like cells. The lung harboured the highest proportions of effector memory-like cells (**Fig 3D**). When we changed the gating strategy by first gating on naïve, memory or effector memory markers and then on Ly6C positivity, we found that about 90% of naïve-like CD27^+^ γδ T cells lack expression of Ly6C across all tissue types (**EV1B**). Similarly, effector memory CD27^+^ γδ T cells were mostly Ly6C^—^. The Ly6C^+^ cells were mainly observed in the memory-like category, making up 40-50% of the population in spleen, LN and lung (**EV1B**). We performed the same analysis on CD4^+^ and CD8^+^ T cells, which showed that only a minority of CD4^+^ T cells expressed Ly6C (**EV1C**). Conversely, CD8^+^ T cells exhibited the same pattern as CD27^+^ γδ T cells, where naïve cells largely failed to express Ly6C and memory/effector memory cells were enriched in Ly6C^+^ cells (**EV1D**). Taken together, these data indicate that for both CD27^+^ γδ T cells and CD8^+^ T cells, Ly6C expression is associated with CD44 expression and memory-/effector memory-like cells in secondary lymphoid organs and peripheral organs.

An important characteristic of CD44^+^ memory T cells is the capability to vigorously proliferate in response to antigen re-exposure (Martin & Badovinac, 2018). Since CD27^+^Ly6C^+^ γδ T cells exhibited a memory-like phenotype (**Fig 3A-C**), we tested the proliferation status of these cells using Ki-67, and we compared this to the proliferation status of CD27^+^Ly6C^—^ γδ T cells in spleen, LN and lung of WT mice. In spleen and LN, CD27^+^Ly6C^+^ γδ T cells showed greater proliferation when compared with CD27^+^Ly6C^—^ γδ T cells (**Fig 3E**). By contrast, both subsets displayed equal proliferative capacity in the lung. Moreover, Ki-67 expression by lung CD27^+^Ly6C^+^ γδ T cells was approximately 6-fold lower than CD27^+^Ly6C^+^ γδ T cells in secondary lymphoid organs. Although it is unclear why the proliferative capacity of lung CD27^+^Ly6C^+^ γδ T cells is different from cells in spleen and LN, these observations support the notion that CD27^+^Ly6C^+^ γδ T cells are proliferative memory-like cells. We then compared these data with the proliferative capacity of CD4^+^ and CD8^+^ T cells from the same mice. Unlike CD27^+^Ly6C^+^ γδ T cells, CD4^+^Ly6C^+^ T cells failed to exhibit greater proliferation in any of the organs examined (**EV1E**). However, CD8^+^Ly6C^+^ T cells followed the same pattern as CD27^+^Ly6C^+^ γδ T cells, where Ki-67 expression was greater in the subset expressing Ly6C in spleen and LN, but not lung (**EV1E**). Taken together, these data further emphasize the similarity between Ly6C-expressing γδ T cells and CD8^+^ T cells.

### γδ T cell subsets proliferate in response to mammary tumours

To determine how the expression of the cytotoxic molecules identified by scRNAseq and the memory-like phenotype of CD27^+^Ly6C^+^ γδ T cells is influenced during cancer progression, we used the *K14-Cre;Brca1*^F/F^*;Trp53*^F/F^ (KB1P) mouse model. This model mimics triple-negative breast cancer, and tumours arising in the mammary glands of these mice are histologically similar to human invasive ductal carcinoma (Liu *et al*, 2007; Millar *et al*, 2020). KB1P mice succumb to mammary tumours at around 25 weeks old. The analysis of tumour-infiltrating γδ T cells was not possible due to the scarcity of these cells at this site. Therefore, we focused our attention on γδ T cells in spleen, LN and lung. First, we measured the proportions of Ly6C-expressing CD27^+^ γδ T cells in age-matched, WT, tumour-free littermates and KB1P tumour-bearing mice. This analysis revealed that CD27^+^ γδ T cells from WT and KB1P tumour-bearing mice express the same levels of Ly6C in spleen, LN and lung (**Fig 4A**). We then compared expression of the cytotoxic molecules identified by scRNAseq (**Fig 1–2**), including CD160, NKG2A, NKp46 and IFNγ, on CD27^+^Ly6C^—^ and CD27^+^Ly6C^+^ γδ T cells in the three different organs. Expression of each molecule remained the same on both CD27^+^Ly6C^—^ and CD27^+^Ly6C^+^ γδ T cells between WT and KB1P tumour-bearing mice with the exception of NKG2A expression on CD27^+^Ly6C^—^ γδ T cells in spleen (**Fig 4B**). These data indicate that mammary tumours in KB1P mice largely fail to influence the expression of these molecules on CD27^+^ γδ T cell subsets. Lastly, we analysed the naïve/memory/effector status between tumour-free and tumour-bearing mice. For CD27^+^Ly6C^—^ γδ T cells, most cells displayed naïve markers (CD62L^+^CD44^—^) in mammary tumour-bearing KB1P mice in a similar fashion to WT mice, regardless of their anatomical location. There was a small increase in the proportion of memory-like CD27^+^Ly6C^—^ γδ T cells in the lung of tumour-bearing KB1P mice compared to WT mice (**Fig 4C**). The vast majority of CD27^+^Ly6C^+^ γδ T cells from WT and tumour-bearing KB1P mice were largely represented in the memory-like group (CD62L^+^CD44^+^). In spleen, the proportion of memory-like CD27^+^Ly6C^+^ γδ T cells was greater in tumour-bearing KB1P mice than in WT mice (**Fig 4C**), suggesting that these cells may have expanded in response to tumour-derived factors. When we measured the proliferative capacity of the γδ T cell subsets, however, Ki-67 expression levels were similar in CD27^+^Ly6C^+^ γδ T cells from WT and KB1P tumour-bearing mice. The CD27^+^Ly6C^—^ γδ T cells from KB1P tumour-bearing mice expressed higher levels of Ki-67 when compared to CD27^+^Ly6C^—^ γδ T cells from WT mice. This observation was consistent across spleen, LN and lung tissue (**Fig 4D**). These data indicate that mammary tumours have little impact on CD27^+^ γδ T cells, but can induce proliferation of CD27^+^Ly6C^—^ γδ T cells through some unknown mechanism without affecting their phenotype.

**Figure 4.**
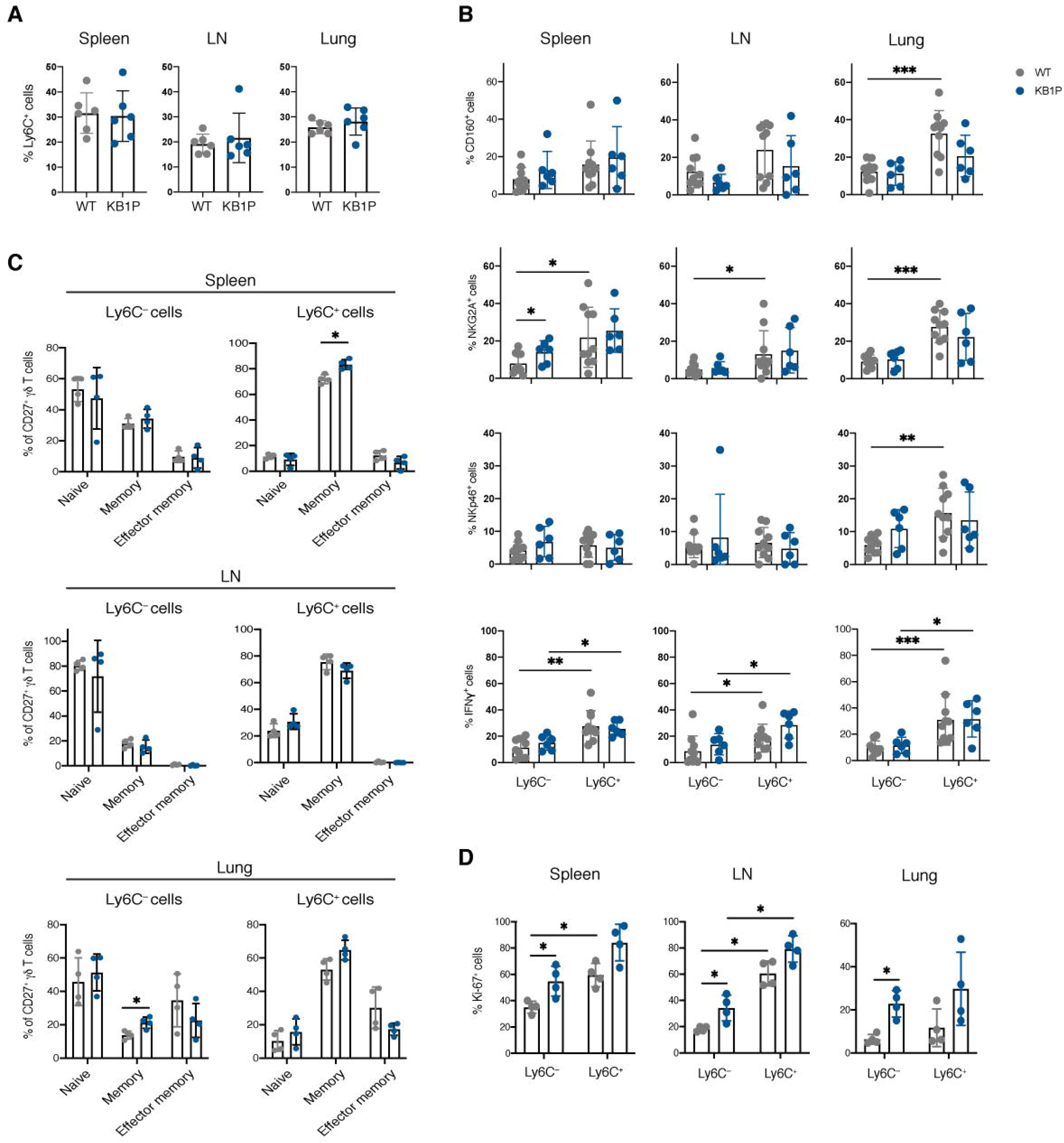
The frequency and phenotype of CD27^+^Ly6C^—^ and CD27^+^Ly6C^+^ γδ T cells are unaffected by mammary tumours. Single-cell suspensions of spleen, LN and lung from WT and *K14-Cre;Brca1*^F/F^*;Trp53*^F/F^ (KB1P) mammary tumour-bearing mice were analyzed by flow cytometry. **(A)** Bar graphs showing the proportion of Ly6C^+^ cells among CD27^+^ γδ T cells in spleen, LN and lung from WT and KB1P mice (n = 6/group). **(B)** Bar graphs showing the proportion of CD160, NKG2A, NKp46 or IFNγ positive cells for CD27^+^Ly6C^—^ and CD27^+^Ly6C^+^ γδ T cell populations in spleen, LN and lung from WT (n = 10/group) and KB1P mice (n = 6/group). **(C)** Bar graphs showing the proportion of naïve, memory and effector memory T cells in CD27^+^Ly6C^—^ and CD27^+^Ly6C^+^ γδ T cells from spleen, LN and lung of WT and KB1P mice (n = 4/group). **(D)** Bar graphs showing the proportion of Ki-67^+^ cells among CD27^+^Ly6C^—^ and CD27^+^Ly6C^+^ γδ T cells from spleen, LN and lung of WT and KB1P mice (n = 4/group). For all panels, each dot represents one mouse. Data are represented as mean ± SD. \**p* < 0.05, \*\**p* < 0.01 and \*\*\**p* < 0.001 as determined by Mann-Whitney U test.

### Depletion of γδ T cells fails to influence mammary tumour growth

Given the proliferative response of CD27^+^Ly6C^—^ and CD27^+^Ly6C^+^ γδ T cells to mammary tumours, we hypothesized that these cells may be reacting to and reactive against the tumour. To test their ability to counteract tumour growth, we first investigated their T cell receptor (TCR) usage to ultimately target the specific TCR with depleting antibodies, since depleting CD27^+^Ly6C^—^ or CD27^+^Ly6C^+^ γδ T cells is not technically possible (**Fig 5A**). This analysis revealed that approximately 60% or more of CD27^+^ γδ T cells from WT mice express the Vγ1 receptor in the three tissues examined, while the other 40% of cells are made up of Vγ4^+^ and Vγ1^—^Vγ4^—^ cells (**Fig 5B**). Next, we examined whether specific TCRs were associated with Ly6C expression. CD27^+^ γδ T cells were analysed in spleen, LN and lung of WT mice as above, followed by gating on Vγ1^+^, Vγ4^+^ or Vγ1^—^Vγ4^—^ cells to determine Ly6C expression on TCR-specific subsets. We found that most (>60%) Vγ1^+^, Vγ4^+^ and Vγ1^—^Vγ4^—^ cells do not express Ly6C regardless of anatomical location (**Fig 5C**), consistent with the overall ratio of Ly6C^—^ to Ly6C^+^ cells (**Fig 2B, 4A**). However, Vγ4^+^ cells expressed higher levels of Ly6C compared to Vγ1^—^Vγ4^—^ cells in spleen and LN (**Fig 5C**). Ly6C expression levels in the lung remained the same for each subset. We then altered the gating strategy to compare TCR usage between CD27^+^Ly6C^—^ versus CD27^+^Ly6C^+^ γδ T cells. Corroborating previous results, both CD27^+^Ly6C^—^ and CD27^+^Ly6C^+^ γδ T cells mainly expressed the Vγ1 TCR chain (**Fig 5D**). In spleen and LN, CD27^+^Ly6C^+^ γδ T cells consisted of more Vγ4^+^ cells and fewer Vγ1^—^Vγ4^—^ cells, than CD27^+^Ly6C^—^ γδ T cells, whereas these subtle differences remained unchanged in the lung (**Fig 5D**).

**Figure 5.**
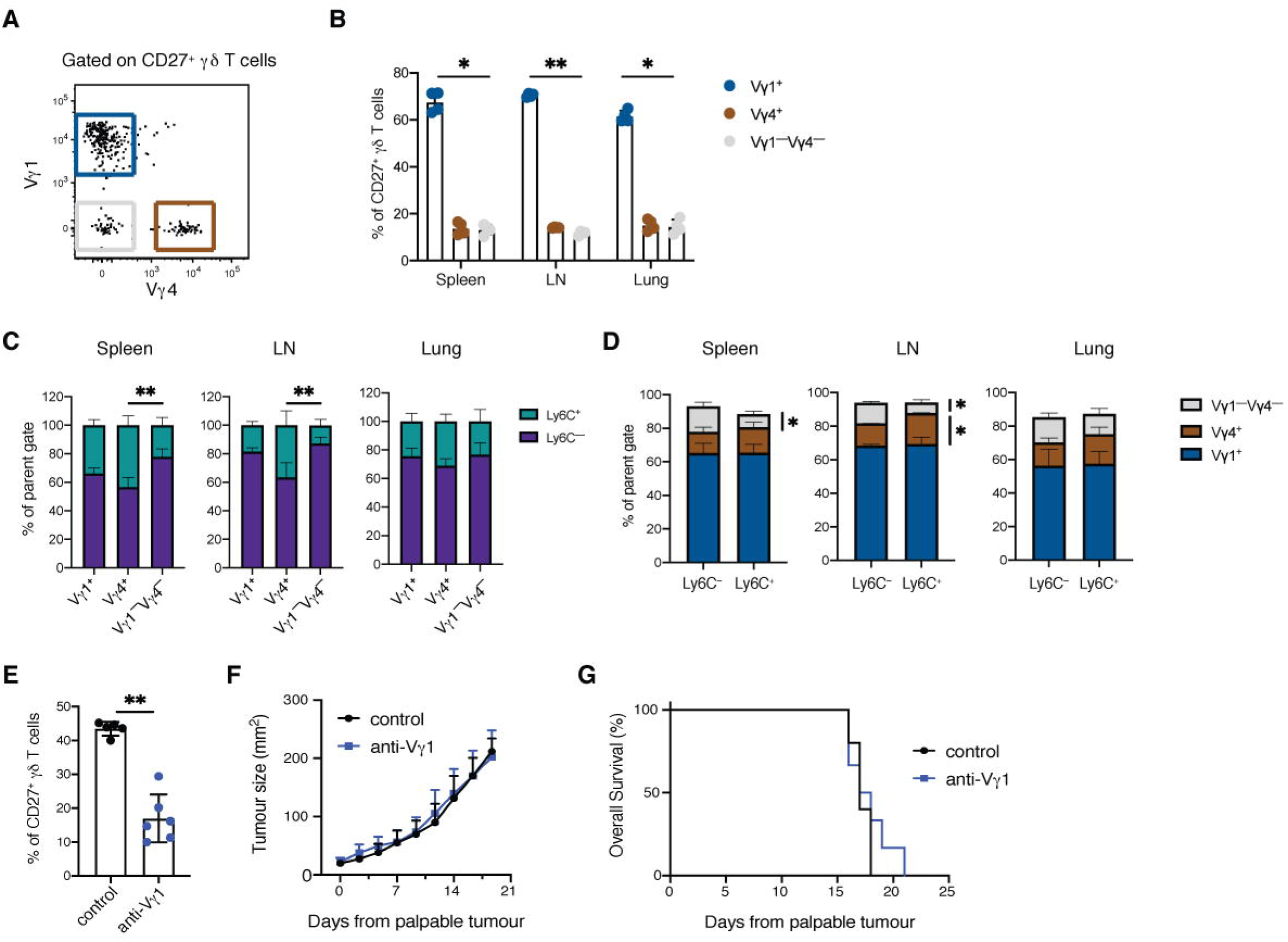
CD27^+^ γδ T cells expressing Vγ1 are dispensable for KB1P tumour growth. T cell receptor (TCR) usage was analyzed on γδ T cells from WT mice by flow cytometry. To assess the function of Vγ1 cells in tumour-bearing mice, mice were transplanted with KB1P tumour fragments; 14 days after tumours were palpable, mice received isotype control antibodies or anti-Vγ1 depleting antibodies daily until tumours reached 15mm. **(A)** Gating strategy for Vγ1^+^ (blue), Vγ4^+^ (brown) and Vγ1^—^Vγ4^—^ (grey) cells after selection of CD3^+^TCRδ^+^CD27^+^ cells. **(B)** Bar graphs showing the proportion of CD27^+^ γδ T cells expressing Vγ1, Vγ4 or Vγ1^—^ Vγδ4^—^ TCRs in spleen, LN and lung (n = 4/group). Data are represented as mean ± SD. \**p* < 0.05 and \*\**p* < 0.01 as determined by repeated measures ANOVA followed by Tukey’s posthoc test. **(C)** Bar graphs showing the ratio of CD27^+^Ly6C^—^ and CD27^+^Ly6C^+^ γδ T cell populations among Vγ1^+^, Vγ4^+^ and Vγ1^—^Vγ4^—^ cells from spleen, LN and lung (n = 4/group). Data are represented as mean ± SD. \*\**p* < 0.01 as determined by repeated measures ANOVA followed by Tukey’s posthoc test. **(D)** Bar graphs showing the ratio of Vγ1^—^Vγ4^—^, Vγ4^+^ and Vγ1^+^ cells among CD27^+^Ly6C^—^ and CD27^+^Ly6C^+^ γδ T cell populations (n = 4/group) from spleen, LN and lung (n = 4/group). Data are represented as mean ± SD. \**p* < 0.05 as determined by Mann-Whitney U test. **(E)** Bar graph showing the proportion of CD27^+^ γδ T cells among CD3^+^ T cells in lungs from mice treated with anti-Vγ1 antibodies (n = 6) or isotype control antibodies (n = 5). Each dot represents one mouse. Data are represented as mean ± SD. \*\**p* < 0.01 as determined by Mann-Whitney U test. **(F)** Tumour growth curve of KB1P mammary tumour-bearing mice treated with anti-Vγ1 antibodies (n = 6 mice) or isotype control antibodies (n = 5 mice). Each dot represents mean ± SD. **(G)** Kaplan-Meier survival analysis of KB1P mammary tumour-bearing mice treated with anti-Vγ1 antibodies (n = 6 mice) or isotype control antibodies (n = 5 mice).

Having confirmed that Vγ1^+^ cells represent the largest proportion of CD27^+^Ly6C^—^ and CD27^+^Ly6C^+^ γδ T cells, we used a Vγ1-depleting antibody to interfere with the endogenous function of these cells in tumour-bearing mice. KB1P tumour fragments were implanted into the mammary glands of syngeneic mice. Once large tumours had formed, mice were treated with isotype control antibodies or anti-Vγ1 antibodies until humane endpoint. Depletion of Vγ1 cells resulted in a 2.5-fold reduction in CD27^+^ γδ T cells (**Fig 5E**). However, their absence failed to affect either primary tumour growth (**Fig 5F**) or survival of KB1P tumour-bearing mice (**Fig 5G**). These data indicate that Vγ1 cells are unable to control late-stage mammary tumours in the KB1P model.

### Expansion of CD27^+^ γδ T cells enhances their cytotoxic phenotype

Previous studies have reported on the anti-tumour functions of CD27^+^ γδ T cells after expansion *in vitro* and adoptive transfer into tumour-bearing mice (Beck *et al.*, 2010; Cao *et al.*, 2016b; He *et al.*, 2010; Liu *et al.*, 2008; Street *et al.*, 2004). Given the inability of endogenous CD27^+^ γδ T cells to control tumour growth in the KB1P tumour transplant model (**Fig 5**), we hypothesized that these cells need further stimulus in order to elicit their cancer-killing functions. To address this hypothesis, we isolated γδ T cells from spleen and LN of WT mice and expanded them *in vitro* with supplementation of the cytokine, IL-15, TCR stimulation through CD3 and co-stimulation through CD28. This expansion protocol lead to a six-fold expansion of CD27^+^ γδ T cells over the course of 4 days (**Fig 6A**). We then investigated whether this culture method changed the phenotype of CD27^+^ γδ T cells by comparing cells from the same mouse, before and after expansion. We profiled the expression of the cytotoxic markers identified from the scRNAseq analysis, including Ly6C, CD160, NKG2A, NKp46 and IFNγ. Additionally, we measured the co-stimulatory molecule, CD28, the cancer cell recognition receptor, NKG2D, and the cytotoxic molecule, granzyme B (GzmB). In this experiment, expression of Ly6C, NKG2A and NGK2D on CD27^+^ γδ T cells remained the same before and after the expansion protocol (**Fig 6B**). By contrast, expression of CD28, CD160, IFNγ and GzmB increased profoundly after culture, whereas NKp46 expression was decreased (**Fig 6B**). Next, we examined these markers on the CD27^+^Ly6C^—^ and CD27^+^Ly6C^+^ cell subsets to determine whether the cytotoxic phenotype of Ly6C^+^ cells was enhanced by the expansion protocol. However, we found that expression of CD28, CD160, NKG2D, NKp46, IFNγ and GzmB was the same between CD27^+^Ly6C^—^ and CD27^+^Ly6C^+^ γδ T cells. NKG2A was the one exception among these markers: its expression was higher on Ly6C^+^ cells than Ly6C^—^ cells (**Fig 6C**). These data indicate that the activation of γδ T cells and their cancer-killing phenotype can be encouraged by *in vitro* expansion with IL-15, TCR stimulation and co-stimulation, but this activation is independent of Ly6C expression.

**Figure 6.**
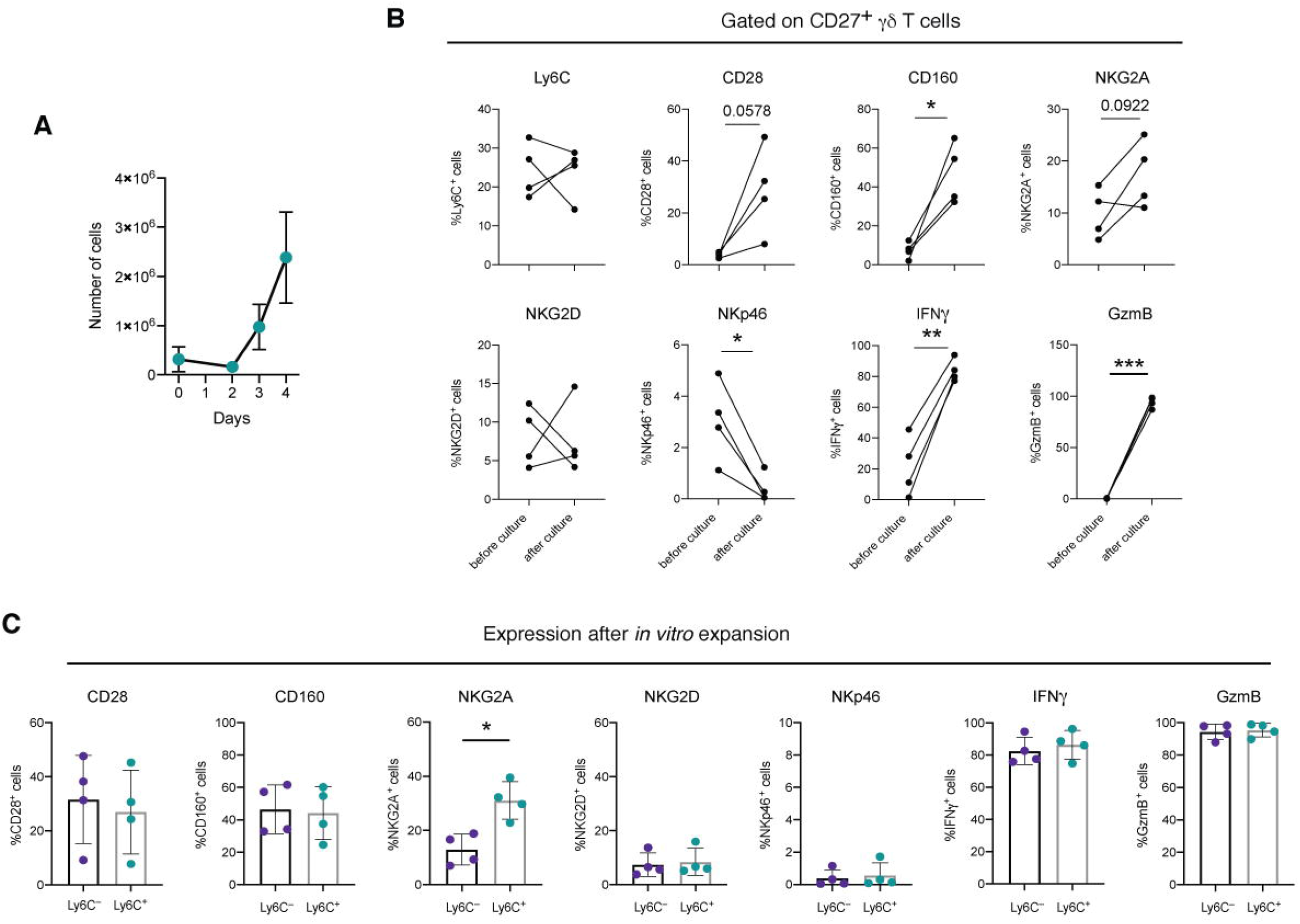
CD27^+^ γδ T cells increase their cytotoxic phenotype after *ex vivo* expansion. CD27^+^ γδ T cells were isolated from LN and spleen of WT mice. The cells were cultured and expanded with CD3/CD28 beads and IL-15. Flow cytometry was used to measure expression of cytotoxic markers before and expansion. **(A)** Line graph depicting the number of CD27^+^ γδ T cells *in vitro* over 4 days (n = 4). Each dot represents mean ± SD. **(B)** Line graphs of the proportion of cytotoxic markers identified by scRNAseq on CD27^+^ γδ T cells before and after expansion (n = 4). Each dot and line represent cells from two pooled mouse samples. \**p* < 0.05, \*\**p* < 0.01 and \*\*\**p* < 0.001 as determined by paired t test. **(C)** Bar graphs of cytotoxic markers expressed by CD27^+^Ly6C^—^ and CD27^+^Ly6C^+^ γδ T cells after expansion *in vitro* (n = 4). Each dot represents cells from one two pooled mice. Data are represented as mean ± SD. \**p* < 0.05 as determined by Mann-Whitney U test.

### *In vitro*-expanded CD27+γδ T cells kill cancer cells

To test the cancer-killing functionality of CD27^+^ γδ T cells before and after expansion, we co-cultured them with two mouse mammary cancer cell lines with different genetic mutations; these cell lines were derived from KB1P tumours or *K14-Cre;Trp53*^F/F^ (KP) tumours. γδ T cells were isolated from spleen and LN of WT mice. Some of these γδ T cells were added directly to KB1P and KP cancer cells, while a proportion of the γδ T cells were expanded with IL-15/CD3/CD28 as above, before co-culture with KB1P and KP cancer cells. Freshly isolated γδ T cells and *in vitro*-expanded γδ T cells were added to mammary cancer cells at the same ratio (10:1). We used the chemotherapeutic agent, cisplatin, to induce cell death as a positive control. After 24 hours, cancer cell death was measured by DAPI uptake. Cisplatin treatment of KB1P and KP cells resulted in a 6-fold and 7-fold increase in cell death, respectively, when compared with untreated cells (**Fig 7A-B**). Freshly isolated CD27^+^ γδ T cells were unable to kill KB1P or KP mammary cancer cells, providing an explanation as to why depletion of Vγ1 cells in KB1P tumour-bearing mice failed in influence cancer progression (**Fig 5**). By contrast, co-culture of *in vitro*-expanded CD27^+^ γδ T cells with KB1P and KP cancer cells achieved a comparable level of cancer cell death as cisplatin treatment. The proportion of dead cancer cells increased by 4-fold in KB1P/γδ T cell co-cultures and 5-fold in KP/γδ T cell co-cultures, when compared to untreated cells (**Fig 7A-B**). To determine whether γδ T cell killing was specific to mammary cancer cells, we also co-cultured *in vitro*-expanded γδ T cells with the lymphoma cell line, YAC-1. Both *in vitro*-expanded γδ T cells and cisplatin effectively induced YAC-1 cell death (**Fig 7C**), suggesting that *in vitro*-expanded γδ T cells have the ability to kill both epithelial- and lymphocyte-derived cancer cells. Taken together, these data indicate that endogenous CD27^+^ γδ T cells can be converted into cytotoxic, cancer-killing cells through *in vitro* stimulation.

**Figure 7.**
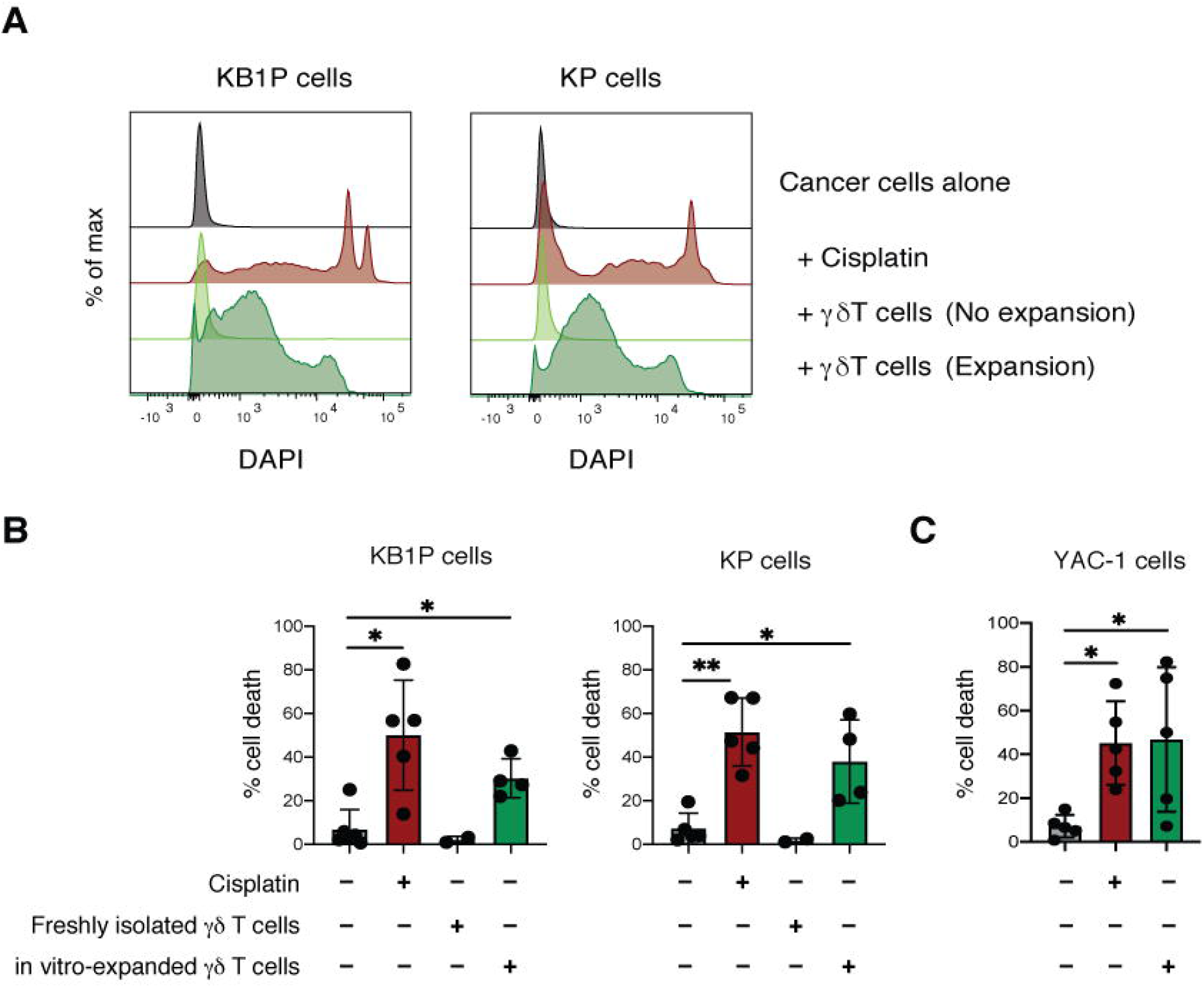
*Ex vivo*-expanded CD27^+^ γδ T cells kill mammary cancer cells. KB1P and KP mouse mammary cancer cell lines were co-cultured with CD27^+^ γδ T cells before and after expansion at a ratio of 1 T cell to 10 cancer cells. Cell death was measured by flow cytometry using DAPI. Cisplatin was used as positive control. **(A)** Representative histograms of DAPI uptake by KB1P or KP mammary cancer cells for four different culture conditions shown. **(B)** Bar graphs showing the proportion of KB1P or KP DAPI^+^ mammary cancer cells (n = 5 control, cisplatin, n = 2 frehsly isolated γδ T cells, n = 4 *in vitro-*expanded γδ T cells). Each dot represents CD27^+^ γδ T cells from two pooled mice. Data are represented as mean ± SD. \**p* < 0.05 and \*\**p* < 0.01 as determined by one-way ANOVA. **(C)** Bar graphs showing the proportion of YAC-1 DAPI^+^ lymphoma cells (n = 5). Each dot represents CD27^+^ γδ T cells from two pooled mice. Data are represented as mean ± SD. \**p* < 0.05 as determined by one-way ANOVA.

## DISCUSSION

While several recent studies report on the heterogeneity of developing γδ T cells in the thymus (Kernfeld *et al*, 2018; Sagar *et al*, 2020; Tan *et al.*, 2019), there is little information on the diversity of CD27^+^ γδ T cells in peripheral organs and secondary lymphoid tissue of adult mice at the single cell level. The data described herein addresses this knowledge gap. Here, we show that CD27^+^ γδ T cells consist of two major transcriptional subsets, defined by naïve T cell markers and cytotoxic, effector-like markers. The transcriptional signatures of each subset were also validated at the protein level. We found that expression of Ly6C – a molecule normally associated with monocytes and neutrophils – stratifies these two subsets and associates with the T cell memory and activation marker, CD44. The phenotype of CD27^+^Ly6C^+^ and CD27^+^Ly6C^—^ γδ T cells and their abundance was relatively similar between the lung and secondary lymphoid organs, suggesting the microenvironmental differences between these tissues have little impact on these cell subsets. However, the lung generally contained more effector memory-like γδ T cells.

Our data support a recent study that labelled a population of γδ T cells expressing *Ccr9* and *S1pr1* mRNA without *Cd44* mRNA in mouse thymus, blood and lymph node as naïve-like cells (Sagar *et al.*, 2020). Although we did not detect *Ccr9* transcript levels in our scRNAseq dataset, the previously identified *Ccr9*-/*S1pr1*-expressing cells are highly analogous to the CD27^+^Ly6C^—^ γδ T cells we describe, which express *S1pr1* mRNA. Our findings corroborate the notion that these thymic-derived cells are immature and expand in secondary lymphoid organs after receiving some unknown stimulus. In addition, these observations raise interesting questions about the relationship between Ly6C^—^ and Ly6C^+^ γδ T cells. Given that *Ly6c2* mRNA did not feature in scRNAseq analyses of thymic γδ T cells (Kernfeld *et al.*, 2018; Sagar *et al.*, 2020; Tan *et al.*, 2019) and the positive association of Ly6C and CD44 on γδ T cells, the data suggest that CD27^+^Ly6C^—^ γδ T cells acquire Ly6C expression together with CD44 following education outside the thymus. Acquisition of Ly6C is presumably accompanied by transcription factors, such as Eomes and T-bet, which regulate expression of cytotoxic molecules (Barros-Martins *et al*, 2016; Lino *et al*, 2017). This potential conversion of Ly6C^—^ γδ T cells into Ly6C^+^ γδ T cells would mirror Ly6C expression on memory CD8^+^ T cells. Indeed, transfer of CD27^+^Ly6C^—^ γδ T cells into lymphopenic mice converts a proportion of these cells into Ly6C^+^CD44^+^ memory-like cells (Lombes *et al.*, 2015). The activation signal that may drive Ly6C^—^ cells into Ly6C^+^ cells remains a mystery, but recent studies suggest that the butyrophilin family, endothelial protein C receptor (EPCR) or other MHC-like molecules are candidate ligands (Willcox & Willcox, 2019). The presentation of these candidate ligands may come from atypical sources, such as mast cells (Mantri & St John, 2019), and may occur outside secondary lymphoid organs where these atypical sources are more abundant.

*Ly6c1* expression is up-regulated in lymph node-derived γδ T cells from old mice, when compared with the same cells from young mice (Chen *et al.*, 2019), suggesting that aging impacts the production of CD27^+^Ly6C^+^ γδ T cells. We found that the frequency of CD27^+^Ly6C^+^ γδ T cells and their phenotype remains unchanged in the KB1P mammary tumour model, when compared to CD27^+^Ly6C^+^ γδ T cells in tumour-free mice. These observations indicate that aging and cancer differentially influence the frequency of CD27^+^Ly6C^+^ γδ T cells. However, it will be important to expand these studies across other mammary tumour models and models of other tumour types to determine whether CD27^+^Ly6C^+^ γδ T cells are affected by tumours with different genetic mutations, especially in tumour-bearing aged mice. From published studies, we know that endogenous γδ T cells counteract tumour growth in several highly immunogenic transplantable models, such as B16 melanoma (Lanca *et al.*, 2013) and methylcholanthrene-induced fibrosarcoma cells (Gao *et al.*, 2003). γδ T cells can also protect against prostate tumour formation in the transgenic mouse model, TRAMP (Liu *et al.*, 2008), as well as spontaneous lymphomas arising in β2-microglobulin/perforin double knockout mice (Street *et al.*, 2004). In addition, it will be important to understand the stage of cancer progression at which CD27^+^Ly6C^+^ γδ T cells may play a role – from tumour initiation to metastasis. In the MMTV-PyMT mammary tumour model, a population of NK1.1/granzyme B-expressing γδ T cells expands in the mammary gland, before epithelial cells transition to the carcinoma stage (Dadi *et al.*, 2016). These γδ T cells have a similar gene expression profile to the CD27^+^Ly6C^+^ γδ T cells we identified (i.e. high levels of *Cd160*, *Klrc1*, *Cd7, Xcl1* mRNA and low levels of *S1pr1* mRNA), suggesting that these two populations are related. Whether the NK1.1/granzyme B-expressing γδ T cells that accumulate in the MMTV-PyMT model during tumour initiation can directly kill cancer cells *in situ* remains unknown.

Future studies are needed to understand the underlying mechanisms by which CD27^+^ γδ T cells kill cancer cells. Our data provides evidence that these innate-like lymphocytes can be imprinted with a memory-like phenotype and can be licensed to attack cancer cells. However, it is unclear which molecules are necessary for cancer cell recognition and induction of cancer cell death. Furthermore, the role of the TCR in this process remains unknown.

The data presented here provide novel information on the degree of homology between mouse and human γδ T cells. Overlaying the gene expression signatures of mouse γδ T cell subsets onto a human PBMC and γδ T cell-enriched dataset revealed that mouse CD27^+^ γδ T cells share a great deal of transcriptional similarity with human circulating γδ T cells. The human dataset contained mature and immature γδ T cells that directly corresponded to CD27^+^Ly6C^+^ and CD27^+^Ly6C^—^ γδ T cells, respectively. Because this dataset included γδ T cells from three individual donors (Pizzolato *et al.*, 2019), we believe that the comparison between species should be done with more human samples, when they become available. Nevertheless, it is clear that the activation status of γδ T cells is conserved across species, providing a greater rationale to use mice for the study of anti-tumorigenic γδ T cells. Our data showing that CD27^+^ γδ T cells acquire the ability to kill mammary cancer and lymphoma cell lines after *ex vivo* expansion corroborate several other studies. MMTV-PyMT mammary cancer cells, 4T1 mammary cancer cells, Lewis lung carcinoma cells, TRAMP-derived prostate cancer cells and B16 melanoma cells die when co-cultured with γδ T cells expanded in the presence of IL-2, IL-15 and/or TCR stimulation (Beck *et al.*, 2010; Cao *et al.*, 2016b; Dadi *et al.*, 2016; He *et al.*, 2010; Liu *et al.*, 2008; Street *et al.*, 2004). Moreover, adoptive transfer of these *ex vivo*-expanded γδ T cells into mice bearing syngeneic tumours delays cancer progression (Beck *et al.*, 2010; Cao *et al*, 2016a; He *et al.*, 2010; Liu *et al.*, 2008; Street *et al.*, 2004). Given the increased interest in exploiting both human Vδ1 and Vγ9Vδ2 cells for cancer immunotherapy (Sebestyen *et al.*, 2020; Silva-Santos *et al.*, 2019), the use of mouse CD27^+^Ly6C^+^ and CD27^+^Ly6C^—^ γδ T cells as surrogates for mature and immature human Vδ1 and Vγ9Vδ2 cells in syngeneic mouse cancer models should overcome several limitations in the field, such as the influence of γδ T cell products on other anti-tumour immune cells in immunocompetent mice. A fully syngeneic platform has considerable potential to improve γδ T cell-based immunotherapies for cancer patients.

## METHODS

### Mice

FVB/n female mice were used in this study. They were purchased from Charles River (6-8 weeks old) or bred from the *K14-Cre;Brca1*^F/F^*;Trp53*^F/F^ (KB1P) colony – a gift from Jos Jonkers lab (Netherlands Cancer Institute). The generation and characterization of KB1P and *K14-Cre;Trp53*^F/F^ (KP) mice has been described previously (Liu *et al.*, 2007). KB1P mice were backcrossed for 4 generations to generate mice at N12. Mice lacking *Cre* recombinase, but containing floxed alleles were used as wild-type (WT) littermates. Mice were born in closed, individually ventilated cages and subsequently moved to open cages. Animals were housed in dedicated barriered facilities on a 12h/12h light/dark cycle and fed and watered *ad libitum*. Animal husbandry and experiments were conducted by trained and licensed individuals. Mice were humanely sacrificed using CO_2_ asphyxiation. Mouse husbandry and experiments were performed in accordance with UK Home Office project licence number 70/8645 (Karen Blyth), carried out in-line with the Animals (Scientific Procedures) Act 1986 and the EU Directive 2010, and sanctioned by local Ethical Review Process (CRUK Beatson Institute and University of Glasgow).

### Single cell RNA sequencing and computational analysis

Lungs were harvested from two 12-16-week-old WT mice and pooled together on two separate occasions. The organs were chopped mechanically with scalpels and transferred into gentleMACS C tubes containing DMEM supplemented with 1μg/mL Collagenase D (Roche) and 25mg/mL DNase I (Sigma-Aldrich). Lung digestion mixes were placed on gentleMACS Octo Dissociator (Miltenyi Biotec) with a heater and digested at 37°C using default programme 37C_m_LDK_01. After dissociation, lung digestion mixes were poured through a 70μm cell strainer with the addition of 2% fetal calf serum (FCS) for enzyme inactivation. Erythrocytes of all single cell suspensions were lysed with ammonium chloride lysis buffer (10X RBC Lysis Buffer, eBioscience). Cells were incubated with Fc receptor block (clone 93; 1:50; Biolegend) for 15 min on ice. Cells were then stained with anti-CD3-FITC (clone 145-2C11; 1:100; eBioscience) and anti-TCRδ (clone GL3; 1:100; Biolegend) for 30 min on ice. After washing in 0.5% BSA in PBS, cells were resuspended in 0.5% BSA in PBS with the addition of DAPI (0.01%) for the exclusion of dead cells. Total live γδ T cells (CD3^+^TCRδ^+^DAPI^—^) were purified by flow cytometry using a FACSAria Fusion (BD Biosciences). Yield was typically 0.0005% of total live lung cells. The sorted cells were loaded onto the Chromium Single Cell 30 Chip Kit v2 (10xGenomics) to generate libraries for scRNAseq (n = 2 mice per experiment). The sequencing-ready library was cleaned up with SPRIselect beads (Beckman Coulter). Quality control of the library was performed prior to sequencing (Qubit, Bioanalyzer, qPCR). Illumina sequencing was performed using NovaSeq S1 by Edinburgh Genomics (University of Edinburgh). The output. bcl2 file was converted to FASTQ format by using cellranger-mkfastq™ algorithm (10xGenomics), and cellranger-count was used to align to the refdata-cellranger-mm10-3.0.0 reference in murine transcriptome and build the final (cell, UMI) expression matrix for each sample. Expression levels were determined and statistically analysed with the R environment (https://www.r-project.org), utilising packages from the Bioconductor data analysis suite (Huber *et al*, 2015) including the SingleCellExperiment (https://bioconductor.org/packages/release/bioc/html/SingleCellExperiment.html">) and Seurat (Stuart *et al*, 2019) packages. After quality control for removal of cells with less than 200 or more than 3000 genes, genes in less than 3 cells and cells with more than 10% from mitochondrial genes, followed by batch correction (Haghverdi *et al*, 2018), we isolated the cells with *Cd27* read values >0. This selection resulted in single cell transcriptomes of 458 CD27^+^ γδ T cells. The first 9 components of the principal component analysis (PCA) were used for unsupervised K-nearest clustering, and non-linear dimensional reduction using t-distributed Stochastic Neighbour Embedding (t-SNE) (Van der Maaten & Hinton, 2008) was utilised for visualization of the data. Then the data were analysed to determine the differentially expressed genes of clusters 0, 1 and 2. Genes from the murine cell clusters 0- and 1-defining signatures were converted to their human orthologs with Ensamble database using the martview interface (http://www.ensembl.org/biomart/martview). Then, by using Single-Cell_Signature_Explorer (Pont *et al.*, 2019), these signatures were scored for each single cell of an already annotated dataset of ~10^4^ human purified γδ T cells and PBMCs (Pizzolato *et al.*, 2019). The signature scores were visualized as heatmap projected on the dataset t-SNE, with contours around those cells scoring > superior quartile.

### Multi-parameter flow cytometry

Lung, spleen and LNs (axillary, brachial, inguinal) were collected from WT mice in phosphate-buffered saline (PBS) and kept on ice until further processing. Single cell suspensions for spleen and LNs were prepared by mashing samples through 70μm cell strainer. Lungs were prepared as above. Cells were placed in V-bottom plates and stimulated with 500X Cell Activation Cocktail with Brefeldin A (Biolegend) in IMDM medium supplemented with 8% FCS, 0.5% β-mercaptoethanol and 100U/mL penicillin/streptomycin for 3 hours at 37°C. Following incubation, Fc receptors were blocked with TruStain FcX (Biolegend) for 20min at 4°C. Antibody cocktails (see Tables) were added for 30min at 4°C in the dark. Dead cells were stained with Zombie NIR Fixable Viability kit (Biolegend) (1:400 in PBS) for 20min at 4°C in the dark. Cells were fixed and permeabilized for intracellular staining with Cytofix/Cytoperm (BD Bioscience), cells were stained with Zombie NIR Fixable Viability kit (Biolegend) (1:400 in PBS) for 20min at 4°C in the dark for dead cell exclusion. Antibodies to intracellular antigens were diluted in permeabilization buffer and incubated for 30min at 4°C in the dark. For intranuclear staining of Ki-67, samples were fixed and permeabilised using FOXP3/Transcription Factor Staining Buffer Set (eBioscience) according to the manufacturer’s instructions. Antibodies used for respective panels are listed in **Table 3**. Samples were acquired using BD LSR II flow cytometer and Diva software. Data analysis was performed by using FlowJo (versions 9.9.6 and 10.7.1) and fluorescence minus one (FMO) controls to facilitate gating. The gating strategy for CD27^+^ γδ T cells is shown in **EV2**.

**Table 3.** Antibodies used for flow cytometry analysis.

| Antigen | Conjugate | Clone | Source | Catalogue | Dilution |
| --- | --- | --- | --- | --- | --- |
| **Cluster validation panel** |  |  |  |  |  |
| **CCR7** | PE/Dazzle594 | 4B12 | Biolegend | 120121 | 1:200 |
| **CD3** | BV650 | 17A2 | Biolegend | 100229 | 1:200 |
| **CD4** | APC-eFluor780 | GK1.5 | Invitrogen | 47-0041-82 | 1:200 |
| **CD8a** | APC-eFluor780 | 53-6.7 | Invitrogen | 47-0081-82 | 1:100 |
| **CD11b** | APC-eFluor780 | M1/70 | eBioscience | 47-0112-82 | 1:800 |
| **CD19** | APC-eFluor780 | 1D3 | eBioscience | 47-0193-82 | 1:400 |
| **CD27** | BV510 | LG.3A10 | BD | 563605 | 1:200 |
| **CD160** | PerCP-Cy5.5 | 7H1 | Biolegend | 143007 | 1:100 |
| **IFNγ** | PE-Cy7 | XMG1.2 | eBioscience | 25-7311-82 | 1:200 |
| **Ly6C** | AF700 | HK1.4 | Biolegend | 128024 | 1:50 |
| **NKG2A/C/E** | BV605 | 20d5 | BD | 564382 | 1:100 |
| **NKp46** | BV421 | 29A1.4 | Biolegend | 137612 | 1:100 |
| **TCRδ** | FITC | GL3 | eBioscience | 11-5711-85 | 1:200 |
| **Zombie NIR** | N/A | N/A | Biolegend | 423106 | 1:400 |
| **Memory phenotype panel** |  |  |  |  |  |
| **CD3** | BV650 | 17A2 | Biolegend | 100229 | 1:200 |
| **CD4** | BV605 | GK1.5 | Biolegend | 100451 | 1:100 |
| **CD8** | BUV805 | 53-6.7 | BD | 564920 | 1:200 |
| **CD11b** | APC-eFluor780 | M1/70 | eBioscience | 47-0112-82 | 1:800 |
| **CD19** | APC-eFluor780 | 1D3 | eBioscience | 47-0193-82 | 1:400 |
| **CD27** | BV510 | LG.3A10 | BD | 563605 | 1:200 |
| **CD44** | PerCP-Cy5.5 | IM7 | Biolegend | 103032 | 1:100 |
| **CD62L** | BUV395 | MEL-14 | BD | 740218 | 1:400 |
| **EpCAM** | APC-eFluor780 | G8.8 | eBioscience | 47-5791-82 | 1:50 |
| **Ly6C** | AF700 | HK1.4 | Biolegend | 128024 | 1:50 |
| **NKG2A/C/E** | BV711 | 20d5 | BD | 740758 | 1:50 |
| **PD-1** | PE-Cy7 | 29F.1A12 | Biolegend | 135215 | 1:100 |
| **TCRδ** | FITC | GL3 | eBioscience | 11-5711-85 | 1:200 |
| **TCR Vγ1** | PE | 2.11 | Biolegend | 141106 | 1:200 |
| **TCR Vγ4** | APC | UC3-10A6 | Biolegend | 137707 | 1:100 |
| **Zombie NIR** | N/A | N/A | Biolegend | 423106 | 1:400 |
| **CD27<sup>+</sup> gdT cell purity test panel** |  |  |  |  |  |
| **CD3** | BV650 | 17A2 | Biolegend | 100229 | 1:200 |
| **CD4** | BV605 | GK1.5 | Biolegend | 100451 | 1:100 |
| **CD8** | BUV805 | 53-6.7 | BD | 564920 | 1:200 |
| **CD27** | PE/Dazzle594 | LG.3A10 | Biolegend | 124228 | 1:400 |
| **TCRδ** | FITC | GL3 | eBioscience | 11-5711-85 | 1:400 |
| **Zombie NIR** | N/A | N/A | Biolegend | 423106 | 1:400 |
| ***In vitro* phenotyping before and after culture** |  |  |  |  |  |
| **CD3** | BV650 | 17A2 | Biolegend | 100229 | 1:200 |
| **CD4** | APC-eFluor780 | GK1.5 | Invitrogen | 47-0041-82 | 1:200 |
| **CD8a** | APC-eFluor780 | 53-6.7 | Invitrogen | 47-0081-82 | 1:100 |
| **CD11b** | APC-eFluor780 | M1/70 | eBioscience | 47-0112-82 | 1:800 |
| **CD19** | APC-eFluor780 | 1D3 | eBioscience | 47-0193-82 | 1:400 |
| **CD27** | BV510 | LG.3A10 | BD | 563605 | 1:200 |
| **CD28** | BV786 | 37.51 | BD | 740859 | 1:100 |
| **CD160** | PerCP-Cy5.5 | 7H1 | Biolegend | 143007 | 1:100 |
| **Granzyme B** | AF647 | GB11 | Biolegend | 515406 | 1:50 |
| **IFNγ** | PE-Cy7 | XMG1.2 | eBioscience | 25-7311-82 | 1:200 |
| **Ly6C** | AF700 | HK1.4 | Biolegend | 128024 | 1:50 |
| **NKG2A/C/E** | BV605 | 20d5 | BD | 564382 | 1:100 |
| **NKG2D** | PE/Dazzle594 | CX5 | Biolegend | 130214 | 1:100 |
| **NKp46** | BV421 | 29A1.4 | Biolegend | 137612 | 1:100 |
| **TCRδ** | FITC | GL3 | eBioscience | 11-5711-85 | 1:200 |
| **Zombie NIR** | N/A | N/A | Biolegend | 423106 | 1:400 |
| **Killing assay** |  |  |  |  |  |
| **CD3** | FITC | 145-2C11 | eBioscience | 11-0031-86 | 1:100 |
| **DAPI** | N/A | N/A | Sigma | D9542 | 1mg/mL |

### Depletion of Vγ1 cells

WT mice were orthotopically transplanted with KB1P tumour pieces as described (Millar *et al.*, 2020). Mice were palpated three times per week for tumour growth. Mice were given antibodies 14 days after tumours were palpable (~3×3 mm). Animals were injected intraperitoneally with a single dose of 200mg anti-Vγ1 (Clone 2.11, Bioxcell, #BE0257) on day 1 followed by individual injections of 100mg on two consecutive days. Control mice followed the same dosage regime with rat-IgG isotype control (Clone 2A3, Bioxcell, #BP0089). Tumour growth was measured by calipers and mice were euthanized when tumours reached 15mm.

### γδ T cell isolation and expansion

Single cell suspensions were generated from pooled lymph nodes (axillary, brachial, inguinal) and spleens of two WT mice as above. γδ T cells were isolated from this suspension using the γδTCR^+^ T cell Isolation Kit (Miltenyi Biotec) according to the manufacturer’s instructions. In brief, CD3^+^ T cells were enriched by negative selection on a LD column and QuadroMACS separator (Miltenyi Biotec), before positive selection of γδ T cells using MS columns with an OctoMacs separator (Miltenyi Biotec). To yield a purer γδ T cell population, positive selection was performed twice. γδ T cells were cultured in IMDM medium supplemented with 10% FCS, 100U/mL penicillin/streptomycin, 2mM glutamine, 50μM 2-Mercaptoethanol (β-ME), 10ng/mL murine interleukin-15 (IL-15) (PeproTech) and CD3/CD28 coated Dynabeads (Thermo Fisher Scientific) at a 1:1 ratio. Cells were kept in 96 U-well plates (Greiner Bio-One) in normoxic incubators at 37°C and propagated at a density of 4×10^4^ cells per well. Purity test after expansion was performed by flow cytometry to confirm exclusion of CD4^+^ and CD8^+^ T cells (typical purity > 90%).

### Co-culture of cancer cell lines and γδ T cells

The generation of mammary tumour cell lines from the KB1P and KP mouse models has been described previously (Millar *et al.*, 2020). Cell lines were maintained in DMEM supplemented with 10% FCS, 100U/mL penicillin/streptomycin and 2mM glutamine. YAC-1 mouse lymphoma cells were gifted from Francesco Colucci (University of Cambridge) and cultured in RPMI medium supplemented with 10% FCS, 100U/mL penicillin/streptomycin and 2mM glutamine. These cell lines were regularly tested for mycoplasma infection. Target cells (KB1P, KP or YAC-1 cells) were seeded in flat-bottom 96 well plates and allowed to rest for at least 6h. Freshly isolated or *ex vivo-*expanded γδ T cells were added at an effector to target ratio of 10:1. Cocultures were kept in IMDM culture medium without supplementation of IL-15 or CD3/CD28 beads and kept in hypoxic incubators at 37°C for 24h. Cisplatin was used as positive control for cancer cell death (75μM for KP cells, 100μM for KB1P and YAC-1 cells). Cells were collected with trypsin, stained with anti-CD3-FITC (eBioscience; clone 145-2C11; 1:100) and DAPI (1μg/mL) was added to samples immediately prior to acquisition. Samples were acquired using BD LSR II flow cytometer and Diva software. Cancer cell death was measured by DAPI uptake.

### Statistical Analysis and Data Visualization

The nonparametric Mann–Whitney U-test was used to compare two groups, while one-way ANOVA followed by Dunn’s post hoc test or repeated measures ANOVA followed by Tukey’s post hoc test was used to compare groups of three or more. Two-way ANOVA with repeated measures was used to analyze tumour growth curves. The log-rank (Mantel–Cox) test was used to analyze Kaplan–Meier survival curves. Sample sizes for each experiment were based on a power calculation and/or previous experience of the mouse models. Analyses and data visualisation were performed using GraphPad Prism (version 8.4.2) and Adobe Illustrator CS5.1 (version 15.1.0).

### Data Accessibility

scRNAseq data: Gene Expression Omnibus GSEXXXXX.

## ACKNOWLEDGEMENTS

We thank Jos Jonkers for the KB1P and KP mice and Francesco Colucci for the YAC-1 cell line. We thank Catherine Winchester and all Coffelt lab members for critical discussion. We would like to thank the Core Services and Advanced Technologies at the Cancer Research UK Beatson Institute (C596/A17196), with particular thanks to the Biological Services and Flow Cytometry Facilities.

## AUTHOR CONTRIBUTIONS

RW, SCE and SBC conceived the idea for the study. RW, SCE, AH, KK, MT, J-JF, AK and S-JR performed the experiments. RW, SCE, AH, KK, MT, J-JF, AK, S-JR, CM and KB analyzed and interpreted the data. RW, SCE and SBC wrote the manuscript with input from all authors.

**Expanded View Figure 1. Naïve, memory and effector memory status of CD4 and CD8 T cells.**

Single-cell suspensions of spleen, lymph node (LN) and lung from wild-type (WT) mice were analyzed by flow cytometry.

(**A**) Bar graphs showing the proportion of naïve T cells (CD62L^+^CD44^—^), memory (CD62L^+^CD44^+^) and effector memory (CD62L^—^CD44^+^), among CD4 and CD8 T cells in spleen, LN and lung (n = 4/group). Each dot represents one mouse. Data are represented as mean ± SD. \**p* < 0.05 and \*\**p* < 0.01 as determined by repeated measures ANOVA followed by Tukey’s posthoc test.

(**B, C, D**) Bar graphs showing the ratio of Ly6C^+^ cells to Ly6C^—^ among CD27^+^ γδ T cells, CD4 T cells and CD8 T cells after gating on naïve, memory or effector memory cells (n = 4/group). Each dot represents one mouse. Data are represented as mean ± SD. \**p* < 0.05 and \*\**p* < 0.01 as determined by Mann-Whitney U test.

(**E**) Bar graphs showing the proportion of Ki-67^+^ cells among CD4 and CD8 T cells from spleen, LN and lung (n = 4/group). Each dot represents one mouse. Data are represented as mean ± SD. \**p* < 0.05 as determined by Mann-Whitney U test or repeated measures ANOVA followed by Tukey’s posthoc test.

**Expanded View Figure 2. Gating strategy for cluster validation.**

Single-cell suspensions of spleen, LN and lung from WT mice were analyzed by flow cytometry to define CD27^+^Ly6C^—^ and CD27^+^Ly6C^+^ γδ T cells. SSC-A = side scatter-area, FSC-A = forward scatter-area.

