## Supplementary figures and images for "Ly6C defines a subset of memory-like CD27^+^ γδ T cells with inducible cancer-killing function"

### Supplemental Figure 1

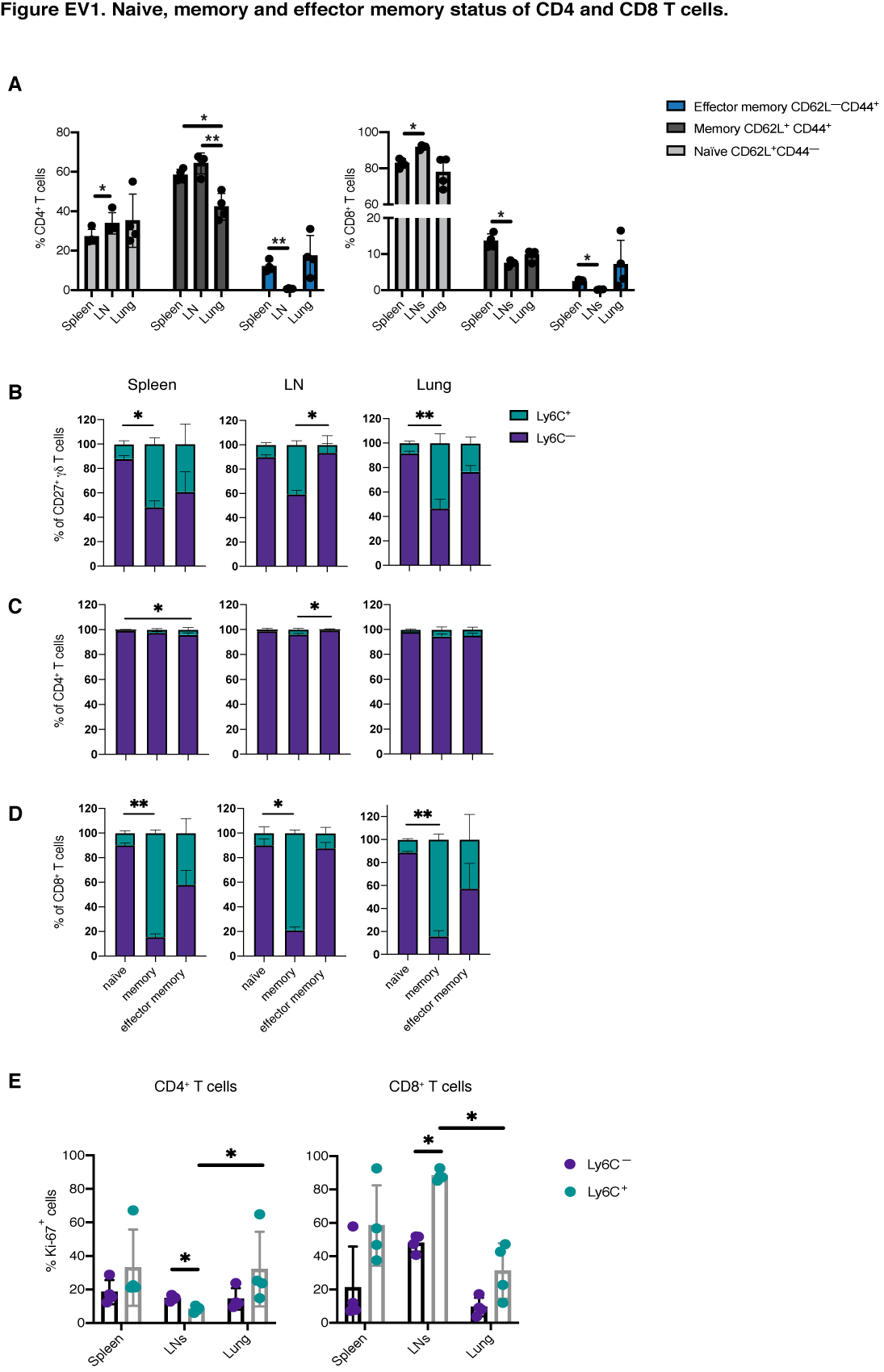

### Supplemental Figure 2

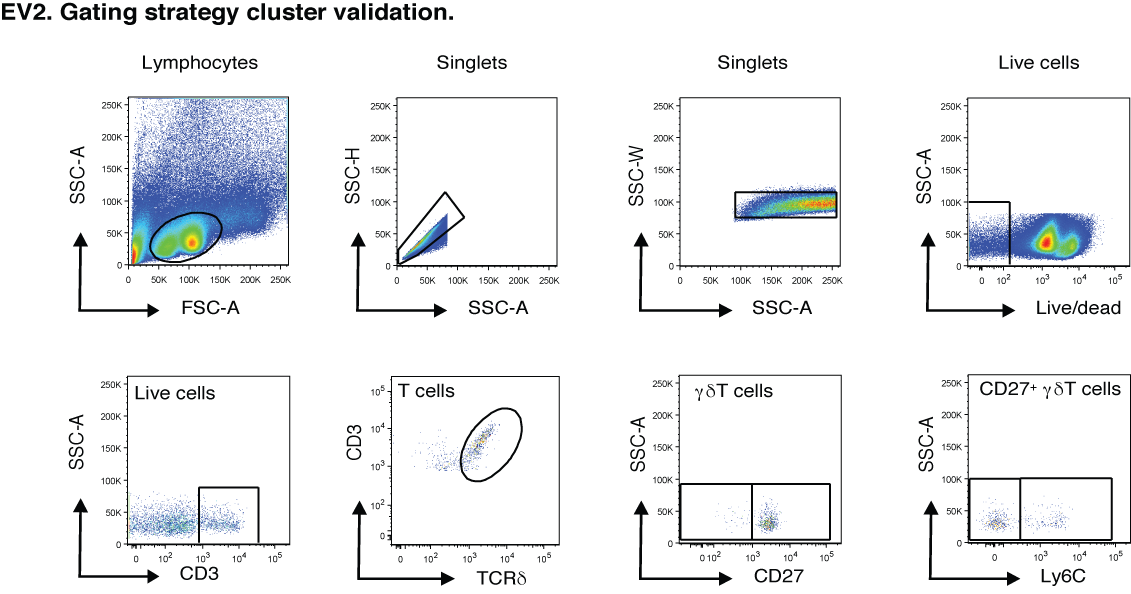
